# Fundamental identifiability limits in molecular epidemiology

**DOI:** 10.1101/2021.01.18.427170

**Authors:** Stilianos Louca, Angela McLaughlin, Ailene MacPherson, Jeffrey B. Joy, Matthew W. Pennell

**Affiliations:** Department of Biology, University of Oregon, USA; Institute of Ecology and Evolution, University of Oregon, USA; British Columbia Centre for Excellence in HIV/AIDS, Vancouver, Canada; Bioinformatics, University of British Columbia, Vancouver, Canada; Biodiversity Research Centre, University of British Columbia, Vancouver, Canada; Department of Zoology, University of British Columbia, Vancouver, Canada; Department of Ecology and Evolutionary Biology, University of Toronto, Toronto, Canada; Department of Medicine, University of British Columbia, Vancouver, Canada

**Keywords:** epidemiology, phylogenetics, statistical inference, birth-death-sampling model

## Abstract

Viral phylogenies provide crucial information on the spread of infectious diseases, and many studies fit mathematical models to phylogenetic data to estimate epidemiological parameters such as the effective reproduction ratio (*R*_*e*_) over time. Such phylodynamic inferences often complement or even substitute for conventional surveillance data, particularly when sampling is poor or delayed. It remains generally unknown, however, how robust phylodynamic epidemiological inferences are, especially when there is uncertainty regarding pathogen prevalence and sampling intensity. Here we use recently developed mathematical techniques to fully characterize the information that can possibly be extracted from serially collected viral phylogenetic data, in the context of the commonly used birth-death-sampling model. We show that for any candidate epidemiological scenario, there exist a myriad of alternative, markedly different and yet plausible “congruent” scenarios that cannot be distinguished using phylogenetic data alone, no matter how large the dataset. In the absence of strong constraints or rate priors across the entire study period, neither maximum-likelihood fitting nor Bayesian inference can reliably reconstruct the true epidemiological dynamics from phylogenetic data alone; rather, estimators can only converge to the “congruence class” of the true dynamics. We propose concrete and feasible strategies for making more robust epidemiological inferences from viral phylogenetic data.

---

For rapidly evolving pathogens, such as RNA viruses, genetic diversity accumulates on the same timescale as transmission [1]. Thus, pathogen genealogies reconstructed from patient samples can provide valuable information on the transmission dynamics of diseases [2, 3]. As sequencing technology and computational methods continue to improve, such phylogenetic approaches [4, 5] are increasingly being used to help inform public health policy during ongoing epidemics, such as during the 2013-2016 Ebolavirus outbreak [6], the 2015-2016 expansion of Zika virus [7], and the SARS-CoV-2 pandemic that began in 2019 [8]. One of the most popular mathematical frameworks used for such phylodynamic inferences is the birth-death (BD) model [9–11], variants of which are also used to reconstruct macroevolutionary dynamics [12]. BD models are typically either fitted to a given time-calibrated phylogeny (henceforth timetree) or jointly estimated with the timetree from molecular sequences, to obtain estimates of the speciation or birth rate (*λ*, corresponding to transmission between hosts in epidemiology or speciation in macroevolution), the extinction or death rate (*µ*, host death or recovery in epidemiology; extinction in macroevolution), and the sampling rate (*ψ*, number of pathogen lineages sampled per time and per extant lineage) through time. From these rates one can calculate critical epidemiological parameters such as the effective reproduction ratio *R*_e_ = *λ/*(*µ* + *ψ*) [9, 10].

Despite the increasing importance of phylodynamic estimates to public health policy, it is generally unknown precisely what information we can hope to extract from phylogenetic data and how robust these estimates are expected to be, particularly when all rates exhibit temporal variation and are unknown a priori. This is important to understand since public policy decisions often need to account for uncertainty in parameter estimates and their implied predictions. Answering this question has become particularly pertinent in light of recent discoveries of major and long unnoticed identifiability issues in BD models in macroevolution, which call into question the reliability of a large number of studies [13, 14]. Specifically, Louca and Pennell showed that if the speciation rate *λ* and extinction rate *µ* vary through time, then there are a vast number of alternative plausible combinations of *λ* and *µ* that could explain any extant timetree (i.e., comprising only extant species) equally well. Such “congruent” scenarios cannot possibly be distinguished from one another statistically, even in the presence of an infinitely large and completely sampled timetree — in other words, it is impossible to design asymptotically consistent estimation methods for *λ* and *µ* based on extant timetrees alone and without strong prior assumptions.

It is largely unknown to what extent and in what form such congruency issues exist in epidemiological BD models. While some relatively minor parameter correlations have been known for special cases [10, 15], an understanding of general identifiability limits is lacking, and the macroevolutionary case has taught us that parameter correlations known for specific cases might severely underestimate the full extent of the problem. This question is non-trivial: while epidemiological BD models appear similar to macroevolutionary BD models, they are more complex because pathogen sequences are typically not sampled at a single final time point. Samples obtained serially through time provide additional information on an epidemic, however new uncertainty is introduced when the sampling rate (*ψ*) is unknown and, especially, when it varies over time.

Here we provide a definite answer to the above questions and demonstrate that, similar to the macroevolutionary case, there are fundamental limits to how much information can be gleaned from timetrees sampled through time in the absence of strong additional constraints. Specifically, we prove mathematically that for any one hypothesized birth-death-sampling (BDS) scenario — i.e., with specific time-varying *λ, µ* and *ψ*, there exists an infinite number of alternative, markedly different and yet plausible BDS scenarios that are statistically indistinguishable from the hypothesized scenario, even with infinitely large phylogenetic datasets. Using simulations and real sequence data from an HIV outbreak in Northern Alberta, Canada we demonstrate that this identifiability issue means that many epidemiological inferences from phylogenetic data alone may not be as well-supported as previously thought. Fortunately, and in contrast to the macroevolutionary case, we are able to identify concrete and feasible approaches towards resolving these issues in practice.

## Identifiability of general birth-death-sampling models

Our starting point is the general birth-death-sampling (BDS) model with arbitrary time-dependent speciation rate *λ*, extinction rate *µ* and sampling rate *ψ*, where we make the common assumptions that sampled lineages (tips in the phylogeny) are immediately removed from the pool of extant lineages and that branching events correspond to transmission events [16]. We use the term “BDS scenario” (or “epidemiological scenario”) to refer to a specific choice of profiles over time for the parameters *λ, µ* and *ψ*. Using mathematical techniques similar to those developed for macroevolutionary models [13], we find that the likelihood of any timetree under a given epidemiological scenario is entirely determined by only two model parameters, called the pulled speciation rate (denoted 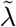) and pulled sampling rate (denoted 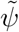). Here 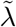 is equal to the speciation rate, *λ*, multiplied by the probability that a lineage is included in the phylogeny, while 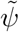 is equal to the sampling rate, *ψ*, divided by the probability that a lineage is included in the phylogeny (overview of symbols in Table 1). The 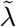 and 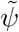 are thus the expected occurrence rate of internal nodes and tips, respectively, over time when divided over the current number of lineages in the tree and in the limit of infinitely large trees (proof in Supplement S.1.3). Note that 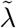 and 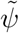 are purely theoretical properties of the BDS scenario that can be calculated from *λ, µ* and *ψ*, and do not depend on any particular data set. We henceforth call any two BDS scenarios congruent if they have the same pulled speciation rate 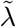 and the same pulled sampling rate 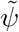. By extension, the congruence class of any BDS scenario henceforth refers to the set of all congruent BDS scenarios. Any two congruent BDS scenarios generate timetrees with the same probability distribution (Supplements S.1.3 and S.1.4), and will yield identical likelihoods for any given timetree. This means that there is no way to distinguish between two congruent scenarios solely based on the properties of sampled timetrees, no matter how large. This result is analogous to that of macroevolutionary BD models (i.e., when all the tips are contemporaneous), where the probability distribution of generated timetrees is entirely determined by the pulled speciation rate, and any two scenarios with identical pulled speciation rates are statistically indistinguishable.

**Table 1:** Overview of main BDS parameters discussed. A parameter is called congruence-invariant if it is identical across congruent BDS scenarios (and thus asymptotically identifiable). Non-congruence-invariant parameters cannot possibly be estimated from phylogenies alone (no matter how large) in the absence of strong additional constraints. Note that each parameter may be time-dependent, and that *τ* denotes age (time before present). Definitions are provided for non-standard BDS parameters.

| symbol | description | definition | congr.-invariant |
| --- | --- | --- | --- |
| $\lambda$ | speciation (transmission) rate | - | no |
| $\mu$ | extinction (recovery+death) rate | - | no |
| $\psi$ | sampling rate | - | no |
| $R_e$ | effective reproduction ratio | $\lambda/(\mu + \psi)$ | no |
| $\delta$ | removal (become uninfected) rate | $\mu + \psi$ | no |
| $S$ | sampling proportion | $\psi/(\mu + \psi)$ | no |
| $E$ | probability of a lineage missing from the phylogeny | Eq. (5) in S.1.1 | no |
| $\tilde{\lambda}$ | pulled speciation rate | $(1 - E) \cdot \lambda$ | yes |
| $\tilde{\psi}$ | pulled sampling rate | $\psi/(1 - E)$ | yes |
| $\tilde{r}$ | pulled diversification rate | $\lambda - \mu - \psi + (1/\lambda)d\lambda/d\tau$ | yes |
| $\tilde{\beta}$ | deterministic branching density | Eq. (18) in S.1.2 | yes |
| $\tilde{\sigma}$ | deterministic sampling density | Eq. (19) in S.1.2 | yes |
| $\tilde{M}$ | normalized deterministic LTT | Eq. (17) in S.1.2 | yes |

Several questions follow from this: For any given BDS scenario, (i) how can one easily determine if another scenario is congruent?; (ii) how many congruent scenarios are there?; and (iii) how different can the epidemiological implications of these congruent scenarios be? To answer these questions, it is useful to consider a number of alternative model parameters, the first of which is called the pulled diversification rate and defined as:

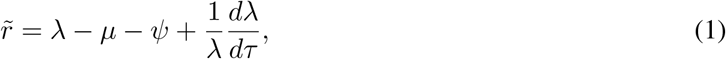

where *τ* denotes age or time before present (Table 1). The pulled diversification rate is equal to the net diversification rate *r* = *λ* − *µ* − *ψ* (or net exponential growth rate in the case of an infectious disease) when the speciation rate *λ* is constant over time, but differs from *r* otherwise. As we prove in Supplement S.1.3, two BDS scenarios are congruent if and only if they have the same pulled diversification rate 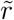 and the same pulled speciation rate 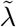. Hence, to check if two BDS scenarios are congruent one can simply compare their 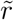 and 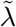. Equivalently, two BDS scenarios are congruent if and only if they exhibit the same deterministic branching density 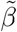 and the same deterministic sampling density 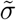, which can be interpreted as rescaled probability densities over time of any randomly chosen observed branching event or sampling event, respectively, in the limit of infinitely large trees (precise definitions in Supplement S.1.2). Hence, the shapes of 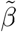 and 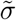 contain easily interpretable information about the temporal distribution of branching and sampling events in the tree. Similarly to 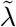 and 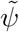, or 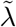 and 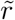, so 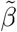 and 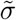 constitute an alternative parameterization of congruence classes, however we stress that all of these parameters do not contain sufficient information for recovering the original model parameters *λ, µ* and *ψ*.

Having introduced the above new parameters, it becomes easy to answer questions (ii) and (iii). For any given scenario (*λ, µ, ψ*) and any given alternative extinction rate *µ*\*, one can find a corresponding *λ*\* and *ψ*\* such that the new scenario (*λ*, µ*, ψ\**) has the same pulled diversification rate and the same pulled speciation rate, i.e., such that the new scenario is congruent to the first one (Supplement S.1.5.2). Indeed, one just needs to solve the differential equation:

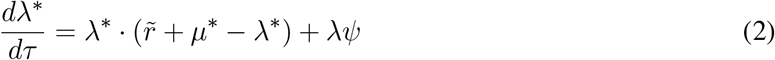

with any initial condition, and choose *ψ*\* = *λψ/λ\**. Many other analogous ways exist for creating congruent scenarios: For example, one can first specify an arbitrary sampling rate *ψ*\* and then adjust *λ*\* and *µ*\* accordingly, or first specify an arbitrary speciation rate *λ*\* and then adjust *ψ*\* and *µ*\* accordingly, or first specify an arbitrary effective reproduction ratio 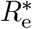 and adjust *λ*, µ\** and *ψ*\* (details in Supplement S.1.5). We note that some scenarios in a congruence class may have negative speciation, extinction, or sampling rates and are therefore biologically irrelevant [17]. Since *µ*\* (or *ψ*\*, or *λ*\*, or 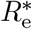) can be chosen nearly arbitrarily and can depend on an arbitrary number of free parameters, the space of congruent BDS scenarios is infinitely large and infinite-dimensional. Many congruent scenarios can appear similarly plausible and similarly complex, and yet exhibit markedly different features, including very different values and opposite trends in *λ, µ, ψ*, or *R*_e_ (examples in Fig. 1).

**Figure 1:**
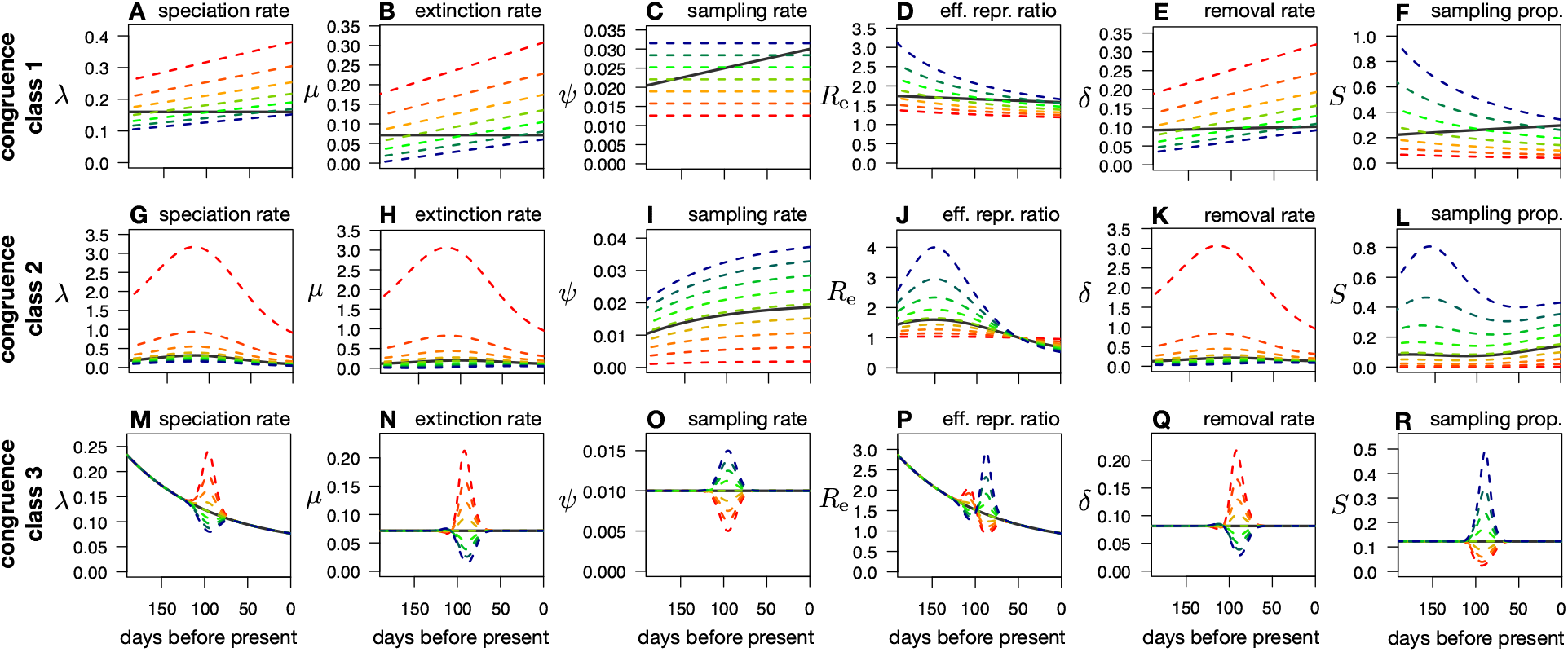
Examples of congruent epidemiological scenarios. (A–F) Transmission (or speciation) rate (A), recovery and death (or extinction) rate (B), sampling rate (C), effective reproduction ratio *R*_e_ = *λ*/(*µ* + *ψ*) (D), removal rate *δ* = *µ*+*ψ* (E), and sampling proportion *S* = *ψ*/(*µ*+*ψ*) (F) of a specific epidemiological scenario (thick black curves), compared to various alternative congruent (i.e., statistically indistinguishable) scenarios (dashed curves). Similarly colored curves across sub-figures correspond to a specific diversification scenario. No viral phylogeny, no matter how large, could possibly distinguish between these (and in fact a myriad of other) scenarios. (G–L) Similar to A–F, but showing scenarios congruent to a different reference scenario. (M–R) Similar to A–F, but showing scenarios congruent to a different reference scenario.

This ambiguity limits the identifiability of epidemiological scenarios when based solely on phylogenetic data, even for infinitely large datasets. Indeed, whatever the true epidemiological history was, there will always exist an infinite number of congruent epidemiological histories. When fitting specific functional forms for *λ, µ* and *ψ*, such as piecewise constant profiles known as skyline models [10], estimators will generally converge to the scenario closest to the congruence class of the true epidemiological history, but not necessarily to the scenario closest to the true epidemiological history itself [17]. Depending on the particular functional forms chosen for *λ, µ* and *ψ*, this can yield markedly different rate profiles over time that may not even approximately resemble the true epidemiological history. This can occur even if the fitted functional forms are flexible enough to adequately approximate the true epidemiological history. In other words, even if one can accurately identify the congruence class that best explains the data (for example, in terms of the 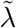 and 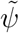, or in terms of the 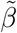 and 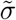), without additional constraints or information one cannot determine which member of the congruence class generated the data. Since the scenarios in a congruent class are indistinguishable for any dataset of any size, with some generally appearing simpler and others more complex than the true epidemiological history, model selection techniques alone cannot resolve this issue (detailed discussion in Supplement S.2). In a Bayesian context, the existence of vastly different congruent scenarios means that the uncertainty in the estimated *λ, µ*, and *ψ* is even more sensitive to the choice of priors than would be apparent from comparing the posterior to the prior distributions of the parameters of some fitted model class. Note that the above issues also apply to other equivalent model parameterizations used in epidemiology, for example based on *R*_e_, the sampling proportion (*S* = *ψ/*(*µ* + *ψ*)) and the removal rate (*δ* = *µ* + *ψ*, also known as “become uninfectious rate”) [10]. We mention that, in contrast, 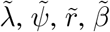 and 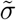 are indeed identifiable given sufficiently large timetrees (Supplement S.1.2).

It was previously demonstrated that for constant-rate scenarios (i.e., where the parameters *λ, µ* and *ψ* do not vary with time) it is impossible to simultaneously estimate *λ, µ* and *ψ* from timetrees alone, because alternative combinations of constant *λ, µ* and *ψ* yield the same likelihood function [10, 11]. It is also known that when fitting BDS skyline models, at least one of the parameters must be fixed in at least one of the time intervals to eliminate correlations between parameter estimates [10]. Our findings are a generalization of these special cases and reveal that these underestimate the true extent of the issue. As soon as one considers that epidemiological parameters could vary arbitrarily over time [18–20], even if they can in principle be approximated by the considered model type (e.g., skyline models), much more information is needed to accurately reconstruct the true epidemiological dynamics.

The above results are analogous to the macroevolutionary case, where the pulled speciation rate is asymptotically identifiable, but does not contain sufficient information for recovering the true speciation and extinction rates. Further, as in the macroevolutionary case, so here two congruent BDS scenarios have identical deterministic lineages-through-time (dLTT) curves (i.e., the number of lineages one would expect in the tree over time based on a deterministic interpretation of the rates *λ, µ* and *ψ*) when conditioned on the number of lineages at some given time point. In contrast to the macroevolutionary case, however, the reverse need no longer be true, i.e., two BDS scenarios with identical dLTTs need not necessarily be congruent. Moreover, while in the macroevolutionary case a congruence class could be uniquely described by a single time-dependent parameter (e.g., the dLTT, or the pulled speciation rate over time), here a BDS congruence class is determined by two time-dependent parameters (e.g., the pulled speciation rate and pulled diversification rate).

### Simulation examples

To demonstrate the challenges for epidemiological inference stemming from the existence of model congruencies, we simulated various hypothetical but realistic epidemiological scenarios and then used two alternative well-established approaches for reconstructing the original dynamics from the generated data. In the first approach, we used the true timetree as an input and estimated the epidemiological dynamics via maximum-likelihood fitting. In the second approach we used nucleotide sequences simulated along the timetree as input, and jointly estimated the timetree together with the epidemiological dynamics using Bayesian MCMC. The latter approach resembles the common situation in molecular epidemiology where the phylogeny is not a priori known, thus introducing additional uncertainty in the reconstruction of the epidemic’s dynamics. To avoid introducing our own biases, for example in the choice of priors, the Bayesian analysis was performed in a blinded way, with some members of our team conducting the simulations and others conducting the Bayesian inference.

For the maximum-likelihood inference we simulated timetrees with >50 000 tips and fitted generic piecewise-linear profiles for *λ, µ* and *ψ* to each timetree, while selecting the optimal number of inflection points using AIC. We used such massive timetrees to avoid errors stemming from small sample sizes, thus focusing on identifiability issues. The fitted models matched the LTTs of the timetrees and the deterministic LTTs of the true scenarios nearly perfectly (Fig. 2D and Supplemental Fig. S1G). The fitted models also adequately explained the timetrees based on 3 different statistical tests performed via parametric bootstrapping [21] (Kolmogorov-Smirnov tests on the distributions of node ages, tip ages, and edge lengths, *P >* 0.05 in all cases). Despite the large sizes of the datasets and the adequacy of the models in explaining the data, the corresponding estimated *λ, µ, ψ, R*_e_, *δ* and *S* were nearly always very different from the true profiles used in the simulations, sometimes even exhibiting opposite trends (Figs. 2A–C and Supplemental Figs. S1A–F and S2A–F). Importantly, the fitted models nearly exactly reproduced the deterministic branching density 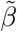 and deterministic sampling density 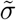 of the true scenario (*R^2^ >* 0.99 in all cases), which, as explained earlier, implies that the fitted models came very close to the true scenario’s congruence class (Figs. 2E,F). This confirms our expectation that fitting yields an estimate of the true epidemiological history’s congruence class, but not necessarily of the true epidemiological history itself. These issues are expected to be even more pronounced with the smaller datasets typical in epidemiology, due to elevated stochasticity.

**Figure 2:**
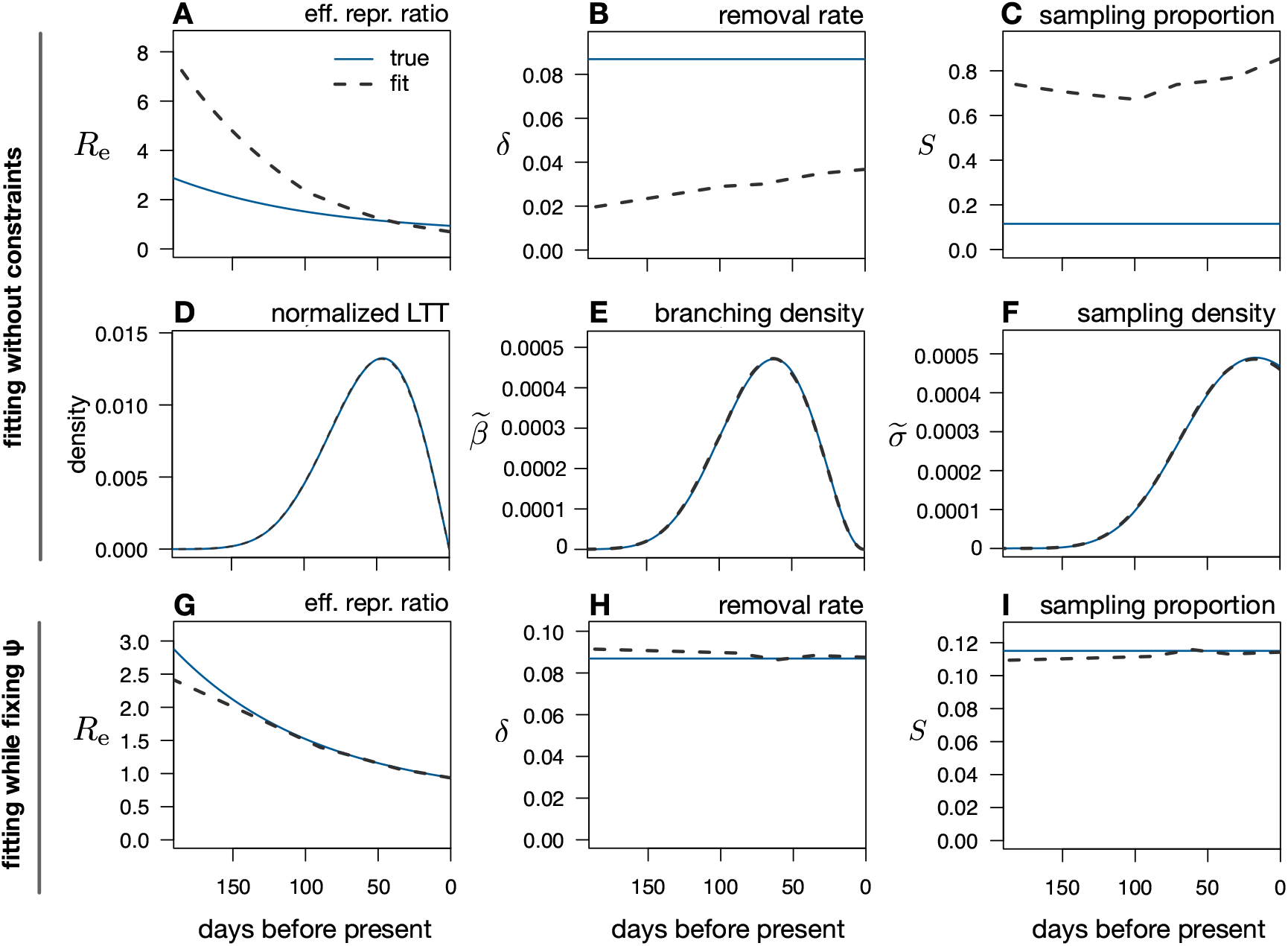
Limits to reconstructing an epidemic’s dynamics via maximum-likelihood. (A–C) Maximum-likelihood estimates (grey dashed curves) of the effective reproduction ratio (*R*_e_), removal rate (*δ* = *µ* + *ψ*), and sampling proportion (*S* = *ψ/*(*µ* + *ψ*)) over time, based on a timetree with 175,440 tips simulated under a hypothetical birth-death-sampling scenario (blue continuous curves). All rates are in day^−1^. Model adequacy was confirmed using predictive posterior simulations with multiple test statistics. Observe the poor agreement between the estimated and true profiles. (D–F) Maximum-likelihood estimates of the deterministic lineages-through-time curve (normalized to have unit area under the curve), deterministic branching density 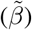, and deterministic sampling density 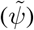, corresponding to the same fitted model as in A–C, compared to their true profiles. The good agreement between the inferred and true profiles shows that the maximum-likelihood fitted model converged towards the true epidemiological history’s congruence class but not the true epidemiological history itself. (G–I) Maximum-likelihood estimates of *R*_e_, *δ*, and *S* inferred from the same data as in A–F, while fixing the sampling rate *ψ* to its true profile. For additional BDS parameters see Supplemental Fig. S1.

For the Bayesian inference we considered two alternative epidemiological scenarios to simulate timetrees of sizes typical in molecular epidemiology (590 and 540 tips, spanning about 15 years, Supplemental Fig. S3). The epidemiological scenarios and the nucleotide substitution models exhibited parameters typically reported for HIV-1 [22, 22–24]. The profiles of *λ, µ, ψ, R*_e_, *δ*, and *S* exhibited moderate variation over time that could be well-approximated using skyline models with three to four intervals. For each of the two scenarios we then conducted a BDS skyline model inference in BEAST2 [5] using the sampled sequences as input data. As indicated above, this inference was internally blinded, i.e., the team members performing the inference had no knowledge of the true epidemiological scenarios used in the simulations. The sole information provided was (a) that model parameters were within the typical ranges known from HIV-1, (b) that four time intervals are sufficient for reasonably approximating the true dynamics with a skyline model (thus avoiding complications in the selection of the number of intervals), (c) the nucleotide substitution model and number of rate categories used, and (d) the value of the present-day sampling proportion, to account for known identifiability issues within skyline models (one parameter in one time interval must be fixed to eliminate correlations between parameters according to Stadler *et al*. [10]). Each of the parameters *R*_e_, *δ*, and *S* varied over time with rate shifts 2, 4, and 8 years before the present, and the sampling proportion in the present was fixed to its known value. The adequacy of the posterior models to explaining the true tree was fully confirmed via predictive posterior simulations [21] based on the same statistical tests as used for the maximum-likelihood fits. The molecular clock rate and the parameters of the nucleotide substitution model were also largely accurately estimated (Supplemental Figs. S10 and S12).

In contrast, in both scenarios the estimated *λ, µ, ψ, R*_e_, *δ*, and *S* deviated substantially from their true profiles, in many cases exhibiting different trends over time or differing by more than an order of magnitude (for example, when considering the maximum-posterior or median-posterior profiles, Fig. 3A–C and Supplemental Figs. S5 and S7), consistent with our expectations. Moreover, the posterior 95% equal-tailed credible intervals barely overlapped with the true profiles of these parameters, suggesting that the inferred posterior distributions severely underestimate the true uncertainty of the parameters. This is not surprising, given that BEAST2 only considered skyline models and does not account for the existence of congruencies. Importantly, in all cases the posterior distributions of the deterministic branching and sampling densities (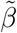 and 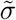) predicted by BEAST2 closely resembled their true profiles (Fig. 3D–F). This implies that BEAST2 accurately inferred the congruence class of the true epidemiological history, just not the true epidemiological history itself (again, consistent with our predictions). To further test this interpretation, we repeated the Bayesian inference while fixing the removal rate to its true profile (approximated by a piecewise constant curve for compatibility with the skyline model). In both scenarios the inferred BDS model parameters much more closely resembled their true profiles (Supplemental Figs. S6 and S8), consistent with the fact that fixing the removal rate profile and the present-day sampling proportion collapses the congruence class to a single scenario (mathematical proof in Supplement S.1.5.7). Together, these results confirm our expectations that any tool attempting to reconstruct an epidemic’s dynamics based on phylogenetic data alone and without strong additional constraints, no matter how large the dataset, can only reconstruct the congruence class of those dynamics rather than the true dynamics.

**Figure 3:**
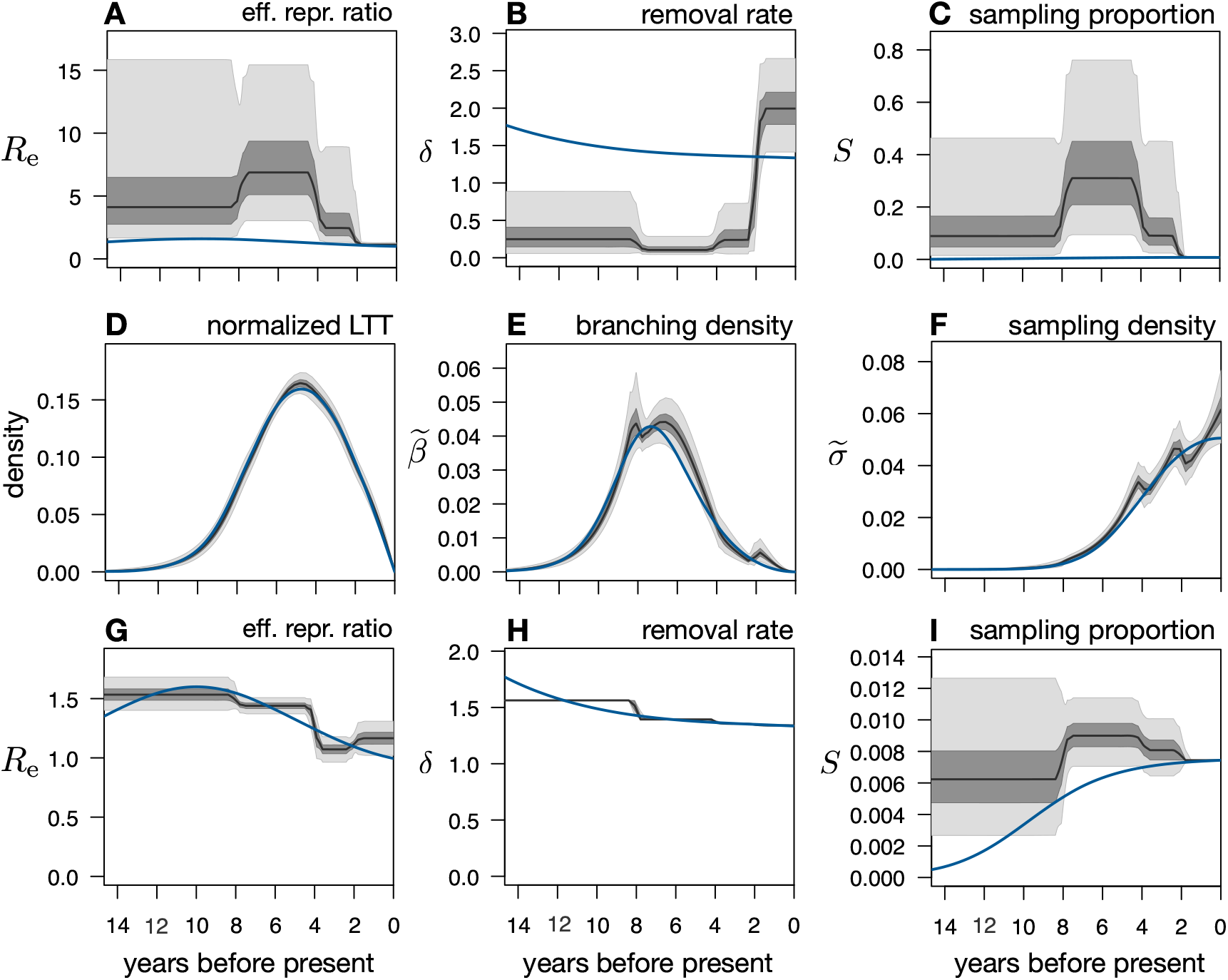
Limits to reconstructing an epidemic’s dynamics in a Bayesian framework. (A–C) Posterior distributions of the effective reproduction ratio (*R*_e_ = *λ/*(*µ* + *ψ*)), removal rate (*δ* = *µ* + *ψ*), and sampling proportion (*S* = *ψ/*(*µ*+*ψ*)), as inferred from 590 sequences simulated under a hypothetical birth-death-sampling scenario (blue curves) using BEAST2. Black curves show posterior median, dark and light shades represent equal-tailed 50%- and 95%-credible intervals of the posterior. All rates are in yr^−1^. The present-day sampling proportion was fixed to its true value during fitting to account for previously reported identifiability issues in skyline models [10]. Model adequacy was confirmed using predictive posterior simulations with multiple test statistics. Observe the poor agreement between the posterior predictions and the true profiles. For additional epidemiological parameters (*λ, µ, ψ*) see Supplemental Fig. S5. For the molecular evolution parameters see Supplemental Fig. S10. (D–F) Distributions of the deterministic lineages-through-time curves (normalized to have unit area under the curve), deterministic branching densities 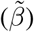, and deterministic sampling densities 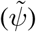, corresponding to the same posterior models as in A–C, compared to their true profiles (blue curves). The relatively good agreement between the inferred and true profiles shows that BEAST2 closely reconstructed the true epidemiological history’s congruence class, but not the true epidemiological history itself. (G–I) Posterior distributions of *R*_e_, *δ*, and *S* inferred from the same data, while fixing the present-day sampling proportion and the removal rate’s profile to their true values. For additional parameters see Supplemental Figs. S6 and S11.

### Illustration for an HIV epidemic

To further illustrate the practical implications of model congruencies, we considered the dynamics of HIV-1 subtype B in Northern Alberta, Canada, over the course of roughly 20 years, reconstructed from 563 molecular sequences using Bayesian BDS skyline models in BEAST2 [5, 10]. We assumed that *R*_e_, *δ*, and *S* varied over time and shifted in 1998 (when triple antiretroviral therapy became available) and in 2010 (to achieve a roughly balanced partitioning of sampling dates). Rate priors were chosen conservatively to reflect the general uncertainty across HIV outbreaks (see Methods) and an uncorrelated log normal relaxed clock was supported over a strict clock based on nested sampling [25]. We found that the posterior distribution of BDS models (Fig. 4E–H) strongly suggests a decline of *R*_e_ over time, a stabilization of the sampling rate in the last two intervals, and a dramatic increase in transmission and recovery rates when comparing the first to the last time interval. The narrow posterior 95% equal-tailed credible intervals for *R*_e_ in the second and third time intervals (Fig. 4H) suggest that *R*_e_ is well-constrained.

**Figure 4:**
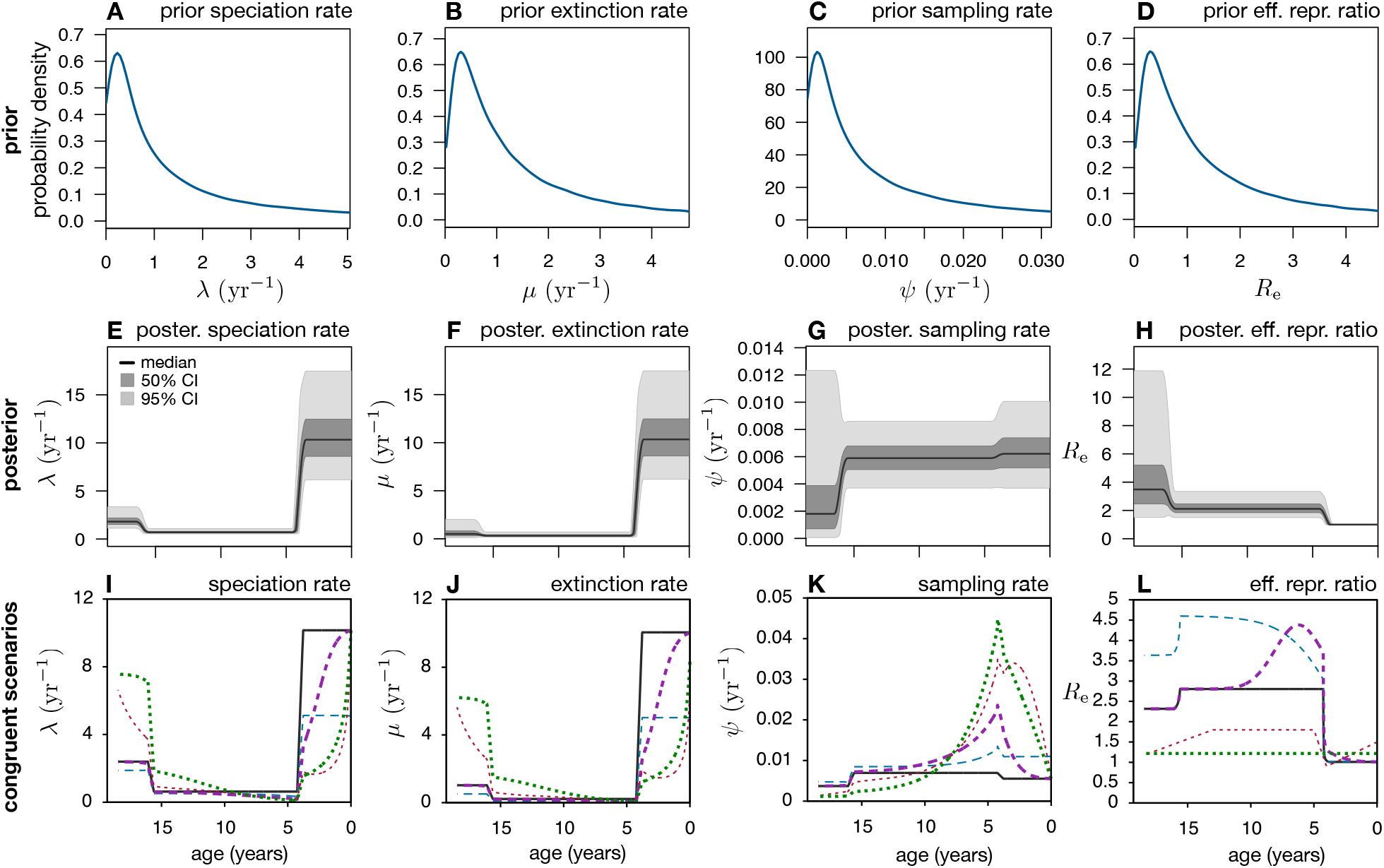
Bayesian reconstruction of HIV spread is compromised by model congruencies. (A–D) Specified prior probability distributions for BDS (skyline) model parameters of HIV-1 subtype B in Northern Alberta, reflecting our a priori knowledge of the plausible range of these parameters. (E–H) Distribution of BDS parameters over time, based on models sampled from the posterior distribution by BEAST2. At each time point, the black curve shows the median value of a parameter across all posterior-sampled models, while the dark and light shadings show 50% and 95% equal-tailed highest posterior density intervals, respectively. (I–L) Maximum posterior probability BDS “reference” scenario (continuous black curves) compared to multiple alternative “congruent” scenarios (dashed curves). Each congruent scenario would generate timetrees with the same probability distribution as the reference scenario, and is thus statistically indistinguishable from the latter. For the posterior distributions of molecular evolution parameters see Supplemental Fig. S19.

However, this posterior is misleading because the inferred credible intervals do not properly capture the ambiguities stemming from model congruencies. Indeed, recall that what we are really estimating is the congruence class of the true epidemiological history, and not the true epidemiological history itself. For illustration, consider the sampled model with maximum posterior probability, shown in Fig. 4I–L. This representative scenario is congruent to a myriad of alternative epidemiological scenarios, many of which are similarly complex and a priori similarly plausible (examples in Fig. 4I–L). In one such congruent scenario, for example, *R*_e_ is constant over time, while in another congruent scenario *R*_e_ temporarily increases. All of these alternative scenarios are equally likely to have generated the data at hand, and this would be true for any other phylogenetic dataset as well. Ruling out some of these congruent scenarios in favor of others requires additional knowledge, such as strong priors on the parameters. Hence, the true uncertainty in the inferred epidemiological parameters is largely determined by the imposed priors, rather than by the computed posterior densities. However, many of the congruent scenarios are not in strong contrast to our priors (Fig. 4A–D), which are typical in the epidemiological literature. In other words, much stronger priors would be needed to collapse the congruence class down to a practical size (e.g., suitably precise for policy decisions), even with massive phylogenetic datasets.

### Ways forward

In our recent analysis of macroevolutionary BD models [13] we proposed that researchers could develop methods to draw insight from asymptotically identifiable variables (i.e., those which are identical between congruent scenarios); in the epidemiological case such quantities include 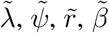, and 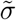. Indeed, for the macroevolutionary case such identifiable variables do contain useful information about historical diversification dynamics [26]. A similar strategy could potentially be fruitful for epidemiological data, although we do not further explore that possibility here. Instead, in the following we discuss a number of ways, some of which build upon current practices in the field, to robustly reconstruct typical epidemiological variables of interest such as *R*_e_.

First, following on what is sometimes done in practice, one can use additional clinical or surveillance data to constrain specific epidemiological parameters. While it is generally recognized that parameter estimation benefits from the use of available constraints, the precise effects of constraints in phylodynamics remained poorly understood and their importance severely underestimated. Our results clarify the amount of information necessary to make an epidemiological scenario asymptotically identifiable. For example, if one of *λ* or *ψ* are known beforehand, then the remaining variables become asymptotically identifiable (details in Supplement S.1.5). This is because in any congruence class there exists at most one scenario with a specific *λ* or *ψ*. Indeed, when we fixed the sampling rate *ψ* to its true profile in our earlier maximum-likelihood fitting tests, the fitted BDS models closely reproduced the true scenarios (Supplemental Figs. S1H–N and S2H–N). Similarly, if *µ* is somehow independently known and either *λ* or *ψ* is known on at least one time point, then the full scenario again becomes asymptotically identifiable. Similar arguments can be made for *R*_e_, *δ*, or *S* (Supplements S.1.5.5, S.1.5.6 and S.1.5.8). Such constraints might be obtained in a variety of ways. For example, in some situations it might be assumed that nearly all people are diagnosed and sampled, in which case the sampling proportion might be fixed to 1. Alternatively, one may estimate the true prevalence of a disease through occasional serological surveys of randomly chosen individuals [27, 28] and then divide the “background rate” of disease detection by that estimate to obtain *ψ*. Further, clinical data may be used to estimate the rate of host death or recovery (i.e., *µ*). During maximum-likelihood fitting, one can fix the independently known parameters. In a Bayesian framework, one can impose appropriate priors to constrain the known parameters. However, we stress that in order to eliminate most congruencies and accurately reconstruct the remaining parameters these priors will need to be much more restrictive than in typical studies [29]. For example, in the second simulation of our Bayesian inference tests described earlier the true removal rate was nearly constant; when we used this information to constrain the removal rate in BEAST2 (i.e., demanding that it is constant across all time-intervals), the parameters *λ, µ, ψ, R*_e_, *δ*, and *S* were estimated much more accurately than in the absence of this constraint (Supplemental Fig. S9). Their accuracy improved further when we fixed the removal rate to its true profile (Supplemental Fig. S8). While constraints such as the above are sometimes included in molecular epidemiological studies, many studies still attempt to estimate the full epidemiological dynamics (*λ, µ* and *ψ*) solely from phylogenies [10, 30, 31]. In contrast, multiple sources of information are commonly utilized when fitting mechanistic epidemiological models, such as differential equation models, or in non-parametric analyses of surveillance data over time. For example, it is common for surveillance data to be combined with independently determined infectivity profiles [32], which are essentially a generalization of *λ*, or with estimates of the serial interval [33], to estimate *R*_e_ over time. Our mathematical results clarify that using additional information beyond just molecular data, e.g., from clinical trials or serological surveys, is not just a means to increase the number of data points — it is essential for ensuring the identifiability of temporally variable epidemiological histories. Developing new phylodynamic models that readily integrate appropriate additional data sources would greatly facilitate this practice.

Second, while molecular epidemiological studies are typically purely observational — i.e., the sampling and data analysis are done independently — this does not need to be the case. Indeed, as we explain here, a properly designed sampling scheme can help reduce identifiability issues. Concretely, short, high-intensity “concentrated” or “contemporaneous” sampling attempts (CSAs), where many individuals are randomly sampled (in addition to sampling symptomatic individuals), can yield valuable information for reconstructing the dynamics of an epidemic and for partly resolving congruencies (details in Supplements S.1.6 and S.1.7). The main requirements are that, first, these CSAs are much shorter than the current transmission and extinction times (i.e., much shorter than 1*/λ* and 1*/µ*), second, *λ* does not differ substantially between the beginning and end of the CSA, and third, the number of lineages sampled during the CSA is much greater than the number of new infections or extinctions occurring in that time period. Such a sampling strategy is not just a hypothetical possibility: during 2020, multiple governments reportedly conducted such CSAs to estimate the seroprevalence of SARS-CoV-2 [34, 35]. If such sampling attempts are performed repeatedly over time and at sufficient temporal resolution, and/or they are combined with other local sequencing data, they can enable an accurate reconstruction of *λ, ψ*, and consequently *µ* over time. For example, when we re-simulated the two hypothetical BDS scenarios used earlier for maximum-likelihood estimation while including 3 CSAs, the subsequently re-fitted models reproduced the true BDS scenarios much more accurately (Supplemental Figs. S1O–U and S2O–U). Note that CSAs differ from the common approach of testing individuals only upon the appearance of symptoms, as in these cases the number of infections detected during any given period tends to be smaller than the number of new infections occurring during that period.

A third potential approach could be to only fit profiles for *λ, µ*, and *ψ* with a strong mechanistic justification, for example derived from deterministic or stochastic differential equation models [16, 36, 37], rather than generic profiles (e.g., skyline models). Whether this approach is effective in avoiding the issues stemming from congruencies in practice is unknown and warrants future investigation. On the one hand, if the true epidemiological history was indeed perfectly described by a given mechanistic model, then fitting that model to a timetree will probably yield accurate parameter estimates, provided of course a sufficiently large dataset. On the other hand, nature rarely (probably never) exactly follows our mechanistic models, and this is certainly true for phenomena with a biological and social component. In this situation, as with generic functional forms, fitting a mechanistic model will generally merely yield a scenario close to the congruence class of the true epidemiological history; in the case of a complex mechanistic model exhibiting a variety of qualitatively different behaviors this might yield a scenario far from the true history itself, even for large datasets.

The above discussion shows that epidemiological reconstruction is generally an easier problem than reconstructing macroevolutionary dynamics. Indeed, the solutions proposed here — experimentally constraining model variables, proper sampling design, and possibly realistic mechanistic models — are currently more tangible in epidemiology than in macroevolution. Well-controlled evolutionary experiments are largely limited to a small number of microorganisms [38, 39], and the fossilization process is outside of a researcher’s control. Further, while detailed mechanistic models (e.g., SIR-type models) are routinely used in epidemiology to analyze surveillance data, in most macroevolutionary studies theoretical models are only conceptual and do not a priori suggest sufficiently precise and accurate functional forms for *λ* or *µ* [40].

We stress that coalescent analysis, an alternative popular framework for reconstructing an epidemic’s effective size (*N*_e_) over time based on pathogen sequences [41, 42], cannot resolve the issues discussed here. While *N*_e_ is in theory asymptotically identifiable provided sufficiently large trees, provided that all assumptions of the coalescent theory are indeed satisfied and provided that the generation time *T* (or serial interval, which is analogous to 1*/λ*) are independently known, the issue of identifiability is essentially being replaced by strong and questionable assumptions about the sampling and transmission process. For example, using solely phylogenetic data coalescent theory can a priori only yield information on the product *N*_e_*T*, and additional assumptions or independent information are needed about *T* in order to actually obtain *N*_e_ and *R*_e_ [43]. In the general situation where *λ* (and hence *T*) can vary arbitrarily through time and is unknown, one is thus faced with similar identifiability issues as with BDS models.

## Conclusions

Our results highlight the limitations of epidemiological inference using phylogenetic data alone. The reported identifiability issues are particularly serious for cases where a reconstruction of historical dynamics is attempted based solely on phylogenetic data and without any additional strong constraints. In such situations, it is generally impossible to reliably reconstruct key epidemiological variables, such as *R*_e_, over time, no matter how large the data. We stress that these issues are separate from the well-recognized errors due to small datasets [36], since any two congruent BDS scenarios remain statistically indistinguishable even for infinitely large phylogenies and hence large phylogenetic datasets alone cannot possibly resolve these issues. Instead, additional data sources beyond just phylogenies, such as from clinical experiments or seroprevalence surveys, are necessary for accurate reconstruction. On a more positive note, we have fully resolved the informational connections between major epidemiological parameters and provide tools for determining their identifiability based on phylogenetic data. In particular, we provide code for exploring the full extent of congruent BDS scenarios and for assessing which epidemiological scenarios can in principle (i.e., with a sufficiently large dataset) be distinguished from one another (Supplement S.3). Our results can help guide proper experimental and sampling designs optimized for epidemiological reconstruction. The solutions discussed also illustrate how the problem of epidemiological reconstruction is rather distinct from that of macroevolutionary reconstruction, in that the former can benefit from additional available sources of information and specific sampling strategies.

## Methods

### Simulations for maximum-likelihood inferences

To demonstrate the implications of model congruencies using simulated trees, we proceeded as follows. Timetrees were generated under two alternative BDS scenarios using the function `generate_tree_hbds` in the R package `castor` v1.6.5 [44] (with options “`include_extant=FALSE, include_extinct=FALSE, no_full_extinction=TRUE`”). In the first scenario, *µ* and *ψ* were assumed to be constant over time while *R*_e_ was exponentially decreasing over time (profiles in Supplemental Fig. S1A–F) and the simulation was halted after 200 days, resulting in a tree with 175,440 tips. In the second scenario, *λ* and *µ* were constant over time while *ψ* increased continuously towards the present (profiles in Supplemental Fig. S2A–F) and the simulation was halted after 200 days, resulting in a tree with 55,934 tips. To each tree, we fitted BDS models with a priori unknown *λ, µ* and *ψ*, each defined as a piecewise linear function of time with inflections at fixed time points (chosen such that their density is approximately proportional to the square root of the tree’s LTT). The optimal number of time-grid points (i.e., the number of fitted parameters for each of *λ, µ* and *ψ*) was chosen by minimizing the AIC of the fitted model [45]. For any given grid size, fitting was done via maximum-likelihood using the `castor` function `fit_hbds_model_on_grid`, with options “`condition=‘auto’, max_start_attempts=100, Ntrials=100`”. Additional epidemiological variables of the fitted models (such as *R*_e_ and the LTT) were computed using the `castor` function `simulate_deterministic_hbds`. To demonstrate the effects of fixing *ψ* to its true value during fitting, we repeated the above fitting process while fixing *ψ* to its true time profile. To demonstrate the effects of concentrated sampling attempts (CSAs), discussed in the main article, we also simulated trees under BDS scenarios similar to the above but modified to include CSAs at 3 discrete time points. We then used these new trees to fit BDS models with unknown *λ, µ* and *ψ*, defined as piecewise linear functions, while also accounting for the added CSAs in the computation of the likelihood [16]. As before, fitting was performed via maximum-likelihood and by choosing the time-grid size according to the AIC. During fitting, the times and intensities (i.e., sampling probabilities) of the CSAs were fixed to their true values (through options “`CSA_ages`” and “`fixed_CSA_probs`”), reflecting general-population randomized seroprevalence surveys in which these properties can be independently determined. The fitted models are shown in Supplemental Figs. S1 and S2. We mention that throughout this article “age” refers to time before present, and “present” refers to the time at which the sampling process was halted.

### Simulations for Bayesian inference

To explore the ability of Bayesian MCMC sampling to recover the true epidemiological history with varying levels of constraints, we conducted an internally-blinded BEAST2 birth-death skyline inference using sampled sequences generated from realistic HIV epidemic simulations. Specifically, group member A simulated timetrees and nucleotide sequences under two different BDS scenarios lasting approximately 15 years, resulting in trees with 590 and 540 tips, respectively (Supplemental Fig. S3). The parameters *λ, µ, ψ, R*_e_, *δ*, and *S* were all within reasonable ranges, i.e., with values within the priors that would typically be specified for an HIV model, with moderate variation through time that can be well-approximated using a skyline model with 3–4 time intervals (Supplemental Figs. S5A–F and S7A–F). Nucleotide sequences of length 1000 bp were simulated along the timetree under an independent-sites HKY substitution model [46] with transition/transversion ratio 5 [24] and stationary base frequencies A:0.4, C:0.17, G:0.21, T:0.22 [22]. The root sequence was chosen randomly according to the stationary base frequencies. Nucleotides were randomly assigned to one of 4 strict-clock substitution rate categories, whose rates were chosen according to a discretized gamma distribution as described by [47], with shape parameter *α* = 0.5 [22–24] and mean rate 2 *×* 10^−3^ yr^−1^.

The simulated sequence alignments and their sampling dates were provided to a second team (“B”) as input data for reconstructing the epidemiological dynamics over time using serial skyline (i.e., piecewise constant) birth-death-sampling models in BEAST2 v2.6.2 with BEAGLE v4.1 [5, 10, 48]. As mentioned earlier, team B did not have knowledge of the true epidemiological parameters, and was initially only provided with the following information: the nucleotide substitution model used (HKY with independent sites), the fact that there were four rate categories according to a discretized gamma distribution, the fact that four time intervals would be sufficient for reasonably approximating the two epidemic histories, the fact that all parameters were chosen within ranges typical for HIV-1, and the present-day sampling proportion. Team B confirmed that there was adequate temporal signal in the sequence data by evaluating the distribution of pairwise patristic distances and divergence over time on a preliminary approximate maximum likelihood tree made in FastTree2 [49] and rooted using residual mean square fitting in Tempest v1.5.3 [50]. Based on the distribution of sampling and branching dates (Supplemental Figs. S3C–F and S4), and to ensure similar amounts of data in each interval, the rate shifts in these models were specified to occur at 2, 4, and 8 years before the present. Skyline models were parameterized in terms of *R*_e_, removal rate *δ*, and sampling proportion *S*, each of which could vary independently in each time interval. In all cases, the present-day sampling proportion was fixed to its true value, to account for known correlations between skyline model parameters [10]. Runs were set up on two independent MCMC chains for 100–200 million states, sampled every 10 000 states (overview of priors in Supplemental Table S1). For each run, log files from both chains were combined using LogCombiner [5] after confirming convergence in Tracer and removing 10% burn-in. We refer to these two runs as 1_444f and 2_444f. For each model drawn from the posterior distribution, we calculated the deterministic LTT, the deterministic branching density, and deterministic sampling density using the castor function `simulate_deterministic_hbds`. Equal-tailed credible intervals of various model parameters were calculated for the posterior distribution of scenarios using the `quantile` function in R. To investigate how the parameter estimates would improve if one were to provide sufficient constraints to collapse the congruence class, we repeated the BEAST2 runs while fixing the removal rate over the entire time period to its true profile (approximated by a piecewise constant curve for compatibility with the skyline model) and while fixing the present-day sampling proportion to its true value (as before); we refer to these new runs as 1_44F4f and 2_44F4f, respectively. Lastly, for the 2nd scenario where the removal rate was nearly constant over time, we investigated how this information might improve parameter estimates, by constraining the removal rate to be constant across all time intervals (with unknown value); we refer to this run as 2_414f. Posterior distributions of epidemiological parameters from runs 1_444f, 1_44F4f, 2_444f, 2_44F4f and 2_414f are shown in Supplemental Figs. S5, S6, S7, S8 and S9, respectively. The corresponding posterior distributions of molecular evolution parameters are shown in Supplemental Figs. S10, S11, S12, S14 and S13. MCMC traces of runs 1_444f and 2_444f are shown in Supplemental Figs. S15 and S16, respectively. The above analyses were also repeated for sequences simulated under a strict molecular clock model with a single discretized rate category from the gamma distribution, yielding similar results.

### Model adequacy tests

To verify that each maximum-likelihood-fitted model was adequate for explaining the timetree, i.e., that parameter estimates were not due to bad model fits, we used parametric bootstrapping to compare various properties of the tree to those expected under the fitted model [21, 51], as follows. For any given tree and fitted model, we simulated 1000 random trees using the model from the root the present-day using the function `generate_tree_hbd_reverse` in the R package castor [52]. We then compared the distribution of tip ages generated by the fitted model to the original tree using a Kolmogorov-Smirnov test. Specifically, for every simulated tree we calculated the empirical cumulative distribution function (CDF) of the tip ages (denoted *F*, and evaluated at the original tree’s tip ages via linear interpolation), and then calculated the average of those CDFs, hence obtaining an estimate for the CDF of tip ages generated by the model (denoted 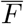). The Kolmogorov-Smirnov (KS) distance between a tree’s CDF *F* and 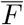, denoted 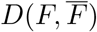, is the maximum distance between *F* and 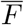 at any age. The statistical significance (*P*) of the original tree’s KS distance 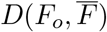 was calculated as the fraction of simulated trees for which 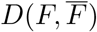 was larger than 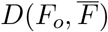. Hence, a small *P* means that the original tree’s distribution of node ages differs substantially from that expected under the fitted model. A similar approach was followed for comparing the model’s and tree’s distribution of node ages or edge lengths. To perform analogous model adequacy tests for our Bayesian analysis, we compared the true timetree (generated during the simulation of the hypothetical epidemic) to the scenarios drawn from the posterior distribution. The methodology was nearly identical to that described above for the maximum-likelihood fits, with the only substantial difference being that each random tree was generated by a scenario randomly chosen from the posterior distribution. The above statistical tests are conveniently implemented in the castor function `model_adequacy_hbds` [44].

### Empirical HIV analysis

For the empirical HIV analysis we used publicly available HIV-1 sequences from Northern Alberta, Canada, collected between 2007 and 2013 and previously described by Vrancken *et al*. [53]. Of the 1055 partial pol sequences (consisting of full *protease* and the first 240 or 400 codons of *reverse transcriptase*) available on Genbank, the analysis was restricted to 809 subtype B sequences, as determined by Vrancken *et al*. based on a maximum likelihood phylogeny of Alberta sequences alongside Los Alamos HIV database sequences (http://hiv.lanl.gov), confirmed using Comet [54]. Four sequences were removed because they were duplicates and one sequence was removed because it had >0.05 ambiguous nucleotides, leaving 804 sequences. Sequences were aligned using mafft v7.402 [55] and known drug resistance mutation sites relative to HXB2 reference were removed [56]. Similarly to Vrancken *et al*., we identified a weak temporal signal within an approximate maximum likelihood tree of all subtype B sequences inferred using FastTree v2.1.11 [49] and rooted by residual mean squared (rms) regression fit in Tempest [50]. The subtype B phylogeny consisted of two deeply split clades herein denoted B.1 (n=624) and B.2 (n=185). After splitting B.1 and B.2 at their MRCA into two trees, re-rooting using rms, and retaining only sequences with residuals < 0.02 substitutions/site (B.1, n=563; B.2, n=164), this yielded an increase in the correlation coefficient of the molecular clock rate fit from 0.21 to 0.33 and 0.34, respectively. Here we focused our analyses on the larger clade (n=563).

The B.1 alignment was used to jointly infer a time-calibrated phylogeny and fit a birth-death skyline serial model in BEAST2 [5, 10]. Model selection consisted of comparing strict and relaxed uncorrelated log normal (UCLN) clocks with free and fixed mean clock rates, as well as multiple rate intervals for *R*_e_, *S*, and */delta* using nested sampling [25]. For all the models compared, site model averaging was conducted using bModelTest [57]; additional priors are summarized in S2. UCLN models with free mean clock rates were more well-supported than their fixed mean clock rate or strict clock model equivalents; and models with rate shifts occurring in 1998 and 2010 had higher likelihoods than their equivalents with equally spaced intervals from origin to the most recent sample. For every model, two parallel MCMC chains of 500 million steps were combined after confirming each run converged, as assessed by effective sample sizes greater than 200 following 10% burn-in. The probability densities of BDS model parameters, based on samples drawn by BEAST from the posterior distribution, are shown in Supplemental Fig. S18. The posterior distributions of molecular evolution parameters are shown in Supplemental Fig. S19. MCMC trace plots are shown in Supplemental Fig. S20.

## Data availability

No raw data were generated for this manuscript. The sequence data from the HIV epidemic in Alberta was described previously Vrancken *et al*. [53] and is publicly available (see Methods for details). The simulated trees and sequences (used for maximum-likelihood fitting and Bayesian analyses) are given in Supplemental File 1.

## Code availability

All analyses were performed using freely available software, indicated throughout the manuscript where appropriate. Novel code for fitting BDS models, examining BDS model congruencies and checking BDS model adequacy (summarized in Supplement S.3) is provided as part of the R package castor, available on The Comprehensive R Archive Network. The BEAST2 configuration XML file for the HIV analysis is provided as Supplemental File 2. The BEAST2 configuration XML files for the analyses of simulated sequences are provided as Supplemental File 3.

## Acknowledgments

We thank Luke Harmon and Josef Uyeda for thoughtful discussions of this work. This project was supported by a GCRC grant from the University of British Columbia. SL and MWP were supported by a US National Science Foundation RAPID grant (#2028986); MWP was also supported by a NSERC Discovery Grant. AML was supported by a CIHR Canada Graduate Scholarships Doctoral award. AM was supported by the EEB department Postdoctoral Fellowship from the University of Toronto. JBJ was supported by a Genome Canada Bioinformatics and Computational Biology grant (287PHY), a Canadian Institutes of Health Research Corona Virus Rapid Response Grant (440371) and the BC Centre for Excellence in HIV/AIDS.

## Author contributions

SL performed the mathematical derivations, numerical simulations and maximum-likelihood fitting, and developed the code for examining BDS model congruencies. AML performed all BEAST analyses and all analyses of the HIV epidemic. All authors contributed to the project’s conception and to the writing of the manuscript.

## Supplementary Information

### S.1 Mathematical derivations

#### S.1.1 General model

The general cladogenic model considered here, and recently formally described by MacPherson *et al*. [16], is a stochastic birth-death-sampling model, where lineages split (“speciate”) stochastically at rate *λ*, disappear (“go extinct”) at rate *µ*, and are sampled (detected) at rate *ψ*. In an epidemiological context, *λ* corresponds to the transmission rate, *µ* to the rate of recovery or death of infected individuals, and *ψ* to the rate at which infected individuals are detected and their pathogen population sequenced. Note that *λ, µ*, and *ψ* can each depend arbitrarily on time. We consider the distribution of time-calibrated reconstructed phylogenies (“time-trees”) obtained through this process, i.e., connecting all sampled variants, after some time *τ*_or_. Note that this cladogenic process is a generalization of the birth-death-skyline model by Stadler *et al*. [10] to the case where *λ, µ*, and *ψ* depend arbitrarily on time rather than in a piecewise constant manner, in the absence of concentrated sampling attempts.

In the following, we count time backward from tips to root, i.e., all variables are expressed as functions of “age” (time before present), with age zero corresponding to the time at which the sampling process is halted. It was recently shown [16] that the likelihood (density) of a given bifurcating timetree generated by a specific BDS scenario starting at age *τ*_or_, conditioned upon the survival and sampling of at least one lineage, is given by:

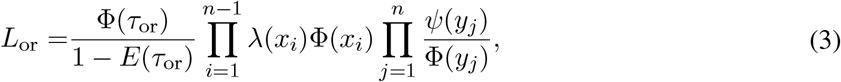

where *n* is the number of tips in the tree, *i* iterates over all branching events (i.e., internal nodes), *j* iterates over all sampling events (i.e., tips), *x*_*i*_ is the age of the *i*-th branching event, *y*_*j*_ is the age of the *j*-th sampling event, *E*(*τ*) is the probability that a lineage alive at age *τ* will not be sampled by the present-day (i.e., is not included in the tree), and Φ is the “flow” of the model [58], defined as:

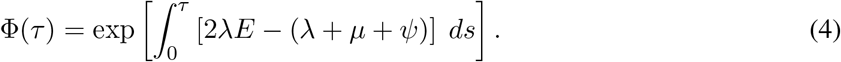

The probability *E* satisfies the differential equation:

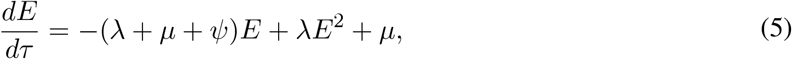

with initial condition *E*(0) = 1. Note that one may alternatively consider the likelihood of the tree conditioned on the age of the tree’s root *τ*_r_, i.e., a splitting occurring at age *τ*_r_ and the eventual sampling of both child lineages:

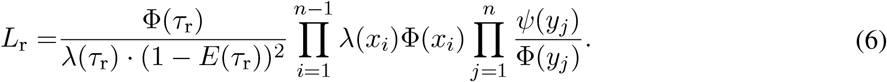

In the limit of infinitely large trees, i.e., when stochastic effects on the LTT curve become negligible, the deterministic LTT curve (dLTT) predicted by the model, denoted *M*, is equal to *M* = *N ·* (1 − *E*), where *M* (*τ*) is the total number of lineages alive at age *τ*. Using the fact that *dN/dt* = (*λ* − *µ* − *ψ*) *· N* as well as Eq. (5), it is straightforward to derive the following differential equation for the model’s dLTT:

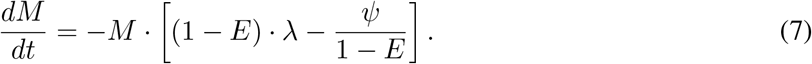

Note that to calculate the dLTT one also needs to specify an initial condition, i.e., condition on the number of lineages at some specific age *τ >* 0 (*M* (0) will always be zero).

#### S.1.2 Pulled parameter

The following composite parameters will be particular useful in our derivations. We define the “pulled speciation rate” 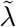 as follows:

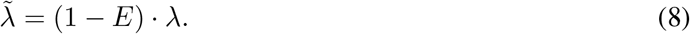

Note that 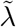 is equal to *λ* if and only if *E* = 0, i.e., if all lineages ever alive are represented in the tree. In the presence of extinction or incomplete sampling, however, 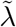 is “pulled” downwards compared to *λ*. Observe that 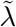 is the expected rate at which internal nodes occur in the tree over time, normalized by the current number of lineages in the tree (*M*) and in the limit of infinitely large trees (see Eq. 7). For sufficiently large trees 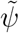 becomes identifiable, since it can just be “read off” the tree’s LTT. We also define the “pulled sampling rate” 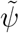 as follows:

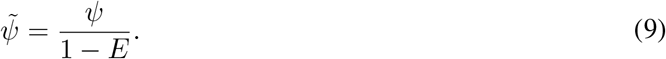

Note that 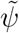 is equal to *ψ* if and only if *E* = 0; otherwise, 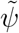 is pulled upward compared to *ψ*. Observe that 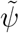 is the expected rate at which tips occur in the tree over time, normalized by the current number of lineages in the tree (*M*) and in the limit of infinitely large trees (see Eq. 7). For sufficiently large trees 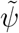 becomes identifiable, since it can just be “read off” the tree’s LTT. We also define the following composite parameter:

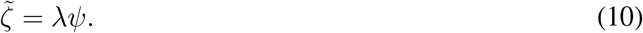

Note that for *τ >* 0 we have 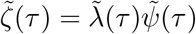. In contrast to *ψ*, which becomes infinite at present day (i.e., at age 0), the parameter 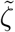 remains finite. Since 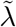 and 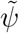 are asymptotically identifiable (i.e., in the limit of infinitely large trees), the same also holds for 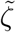. We also define the “pulled diversification rate” 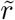 as follows:

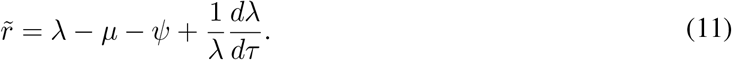

Note that 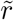 is equal to the net diversification rate *r* = *λ* − *µ* − *ψ* only if *λ* is constant over time. If *λ* varies over time, 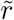 is pulled upwards or downwards compared to *r*, depending on how *λ* varies with time. Taking the derivative on both sides of Eq. (8) and using Eqs. (5) and (10) it is straightforward to show that 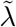 satisfies the differential equation:

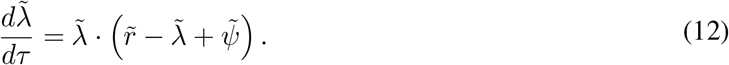

Note that we can also write Eq. (12) as:

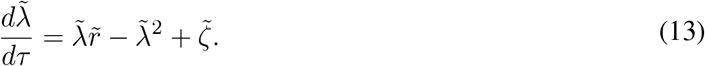

Hence, 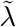 is fully determined by 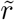 and 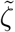 (note that 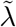 always satisfies the initial condition 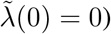(0) = 0). Solving Eq. (12) for 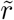 yields:

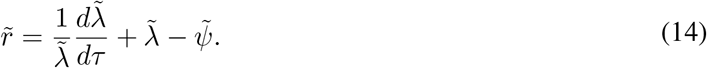

Since 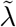 and 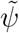 are asymptotically identifiable, the same also holds for 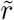. Solving Eq. (12) for 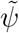 yields:

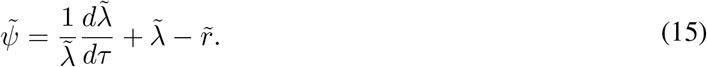

Thus, 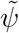 is fully determined by 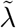 and 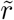. Multiplying Eq. (15) by 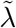 yields:

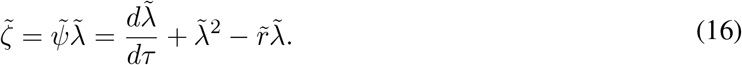

Hence, 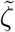 is fully determined by 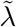 and 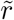. It is also worth introducing three additional parameters, the “normalized dLTT”:

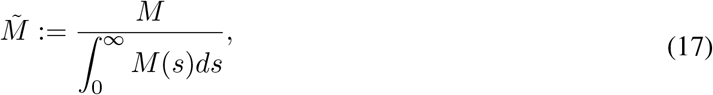

the “deterministic branching density”:

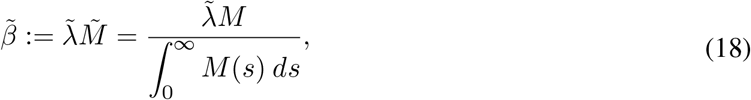

and the “deterministic sampling density”:

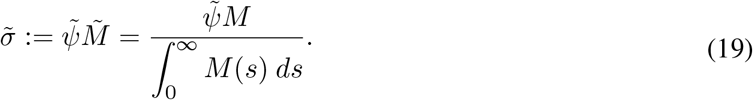

Note that neither 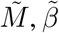 nor 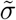 depend on the specific dLTT, as long as the latter solves the differential equation (7), in other words 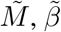 and 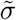 are independent of the specific initial condition chosen for *M*. Hence, 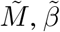 and 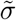 are fully determined by 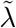 and 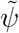, and are thus asymptotically identifiable. Reciprocally, 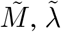, and 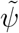 are fully determined by 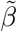 and 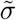. Indeed, from Eq. (7) we have:

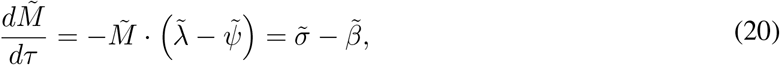

with initial condition *X*(0) = 0. Hence:

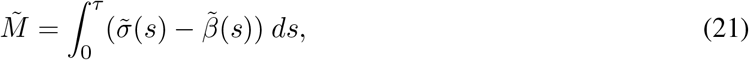

and:

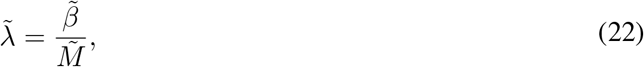

and:

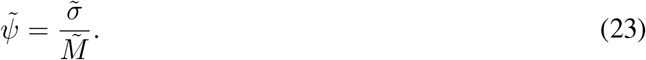

The normalized dLTT is simply the dLTT rescaled so that its area-under-the-curve is equal to 1; hence, the normalized dLTT is a property of the particular BDS scenario that does not depend on the specific initial condition for *M*, and which can be interpreted as a probability density over time. The numerator 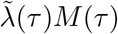 on the right-hand-side of Eq. (18) is the expected rate of branching events observed in the tree at age *τ*, while the denominator 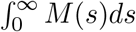 merely acts as a rescaling. Hence the normalized version of 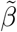 (i.e., if divided by its integral 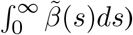 can be interpreted as the probability density on [0, ∞) for any randomly chosen observed branching event. The normalization of 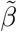, however, is such that the integral 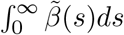 is equal to the average pulled speciation rate, where the average is calculated based on the normalized dLTT:

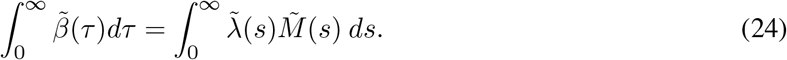

In particular, 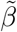 contains information not just about the shape of the branching rate over time, but also about the absolute magnitude of the pulled speciation rate. A similar interpretation exists for the deterministic sampling density 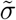, i.e, 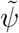 is the (non-normalized) probability density on [0, *∞*) for any randomly chosen observed tip. In contrast to 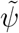, the sampling density 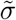 stays finite at age 0:

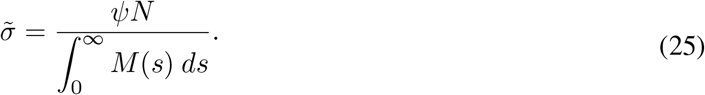

Note that the interpretations of some of the above parameters are strictly speaking only valid in the limit of infinitely large trees, however the parameters themselves are simply properties of the BDS scenario and thus well-defined regardless of the size of the tree.

#### S.1.3 Congruent models

From Eq. (3) and Eq. (6) it becomes clear that the likelihood of a timetree only depends on the sampling and branching ages in the tree, but not the precise tree topology itself. As we show below, a similar statement can also be made about the model’s role, as follows: Any two birth-death-sampling scenarios with the same pulled speciation rate 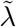 and the same pulled sampling rate 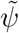 will always yield the same likelihoods. We thus henceforth call two birth-death-sampling scenarios “congruent” if they have the same 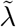 and the same 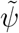. Equivalently, following Supplement S.1.2, two scenarios are congruent if and only if they have the same 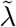 and the same 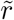, or equivalently, if and only if they have the same 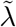 and the same 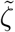, or equivalently, if and only if they have the same 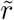 and the same 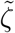, or equivalently, if and only if they have the same deterministic branching density 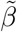 and the same deterministic sampling density 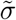

To prove that congruent BDS scenarios have the same likelihoods regardless of the data, we proceed as follows. From Eq. (4) we have:

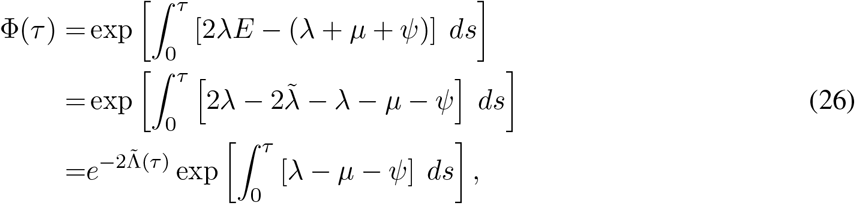

where we defined:

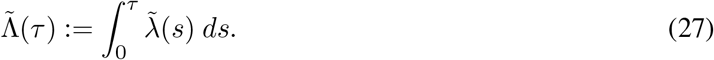

Using Eq. (11) we can further write:

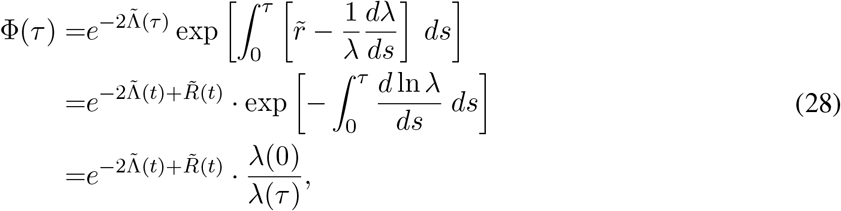

where we defined:

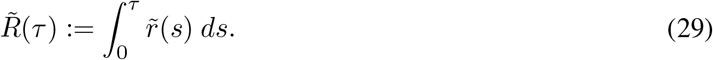

Using Eq. (28) in the formula for the likelihood, Eq. (3), yields:

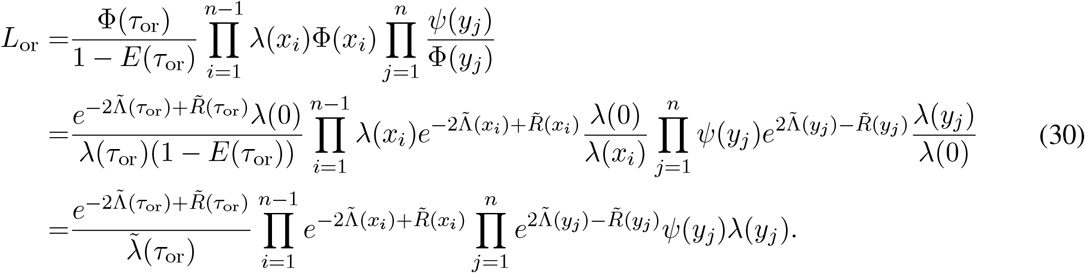

Since 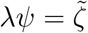, we can further write:

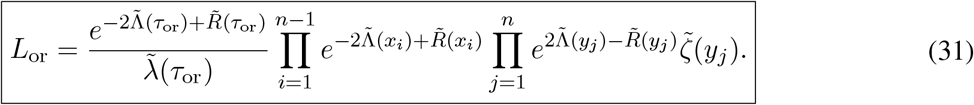

Note that the likelihood in Eq. (31) depends solely on the variables 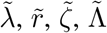 and 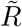, all of which are fully determined by the pulled speciation rate 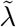 and pulled sampling rate 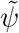

Using a similar approach one can also write the likelihood *L*_r_, i.e. conditioned on the root age, in a format that only depends on pulled variables:

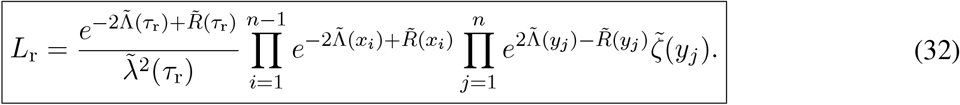

### S.1.4 Congruent models have identical distributions of tree sizes

In the following we show that any two congruent BDS scenarios will generate trees whose size (i.e., number of tips) has the same probability distributions, when conditioned on the age of the stem and the sampling of at least one tip (or the age of the root, and the sampling of both daughter lineages). Let *P*_*n*_(*τ*) be the probability that a lineage alive at age *τ* will have exactly *n* sampled descendants in the final tree (i.e., at present-day, where the process ends). It is straightforward to show that *P*_*n*_ (for any *n ∈ {*0, 1, 2, .. *}*) satisfies the differential equation:

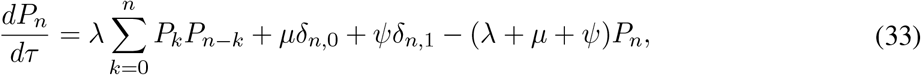

with initial condition:

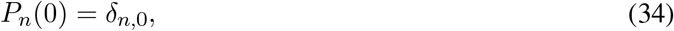

where *τ* denotes age and *δ* denotes the Kronecker-delta symbol, i.e. *δ*_*n,k*_ is 1 if *k* = *n* and 0 otherwise. Denote 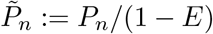. We wish to show that 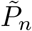 is the same for any two congruent BDS scenarios, for any *n >* 0. We have:

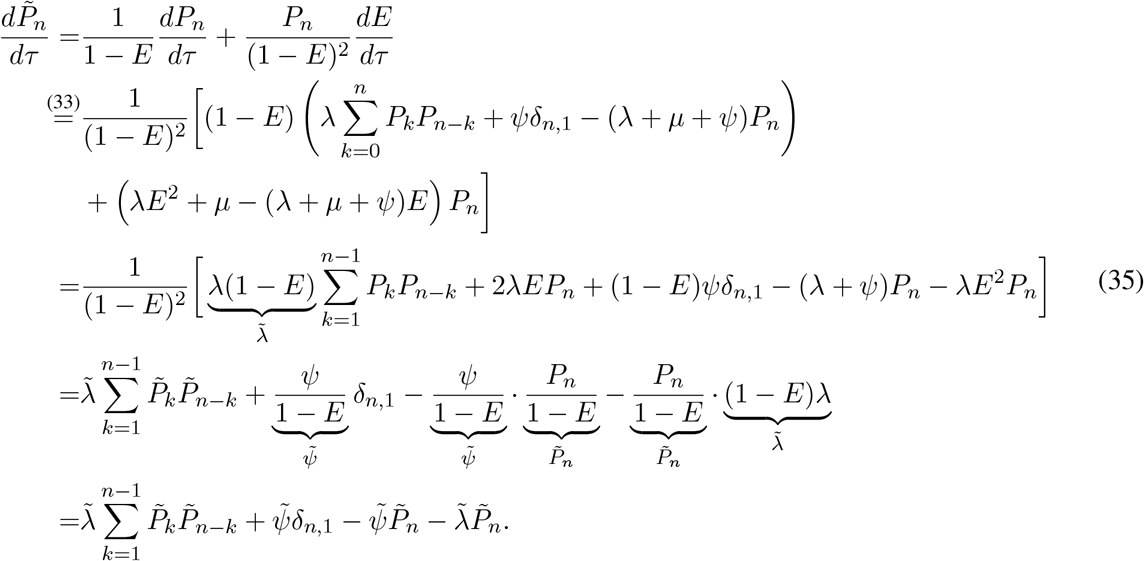

Further, according to L’Hopital’s rule one has:

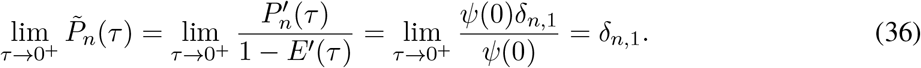

Since any two congruent scenarios have the same 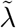 and the same 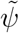, each of their 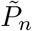 (with *n ≥* 1) will satisfy the same differential equation (35) with the same initial condition (36). We thus conclude that any two congruent scenarios will have the same conditional probabilities 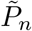.

A similar conclusion can be drawn for the case where we condition on the age of the root and the survival of its two child lineages, as follows. Denote by 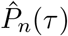 the conditional probability that the tree has size *n*, given that the root split at age *τ* and that both of its child lineages survived. Then for any *n ≥* 2 we have:

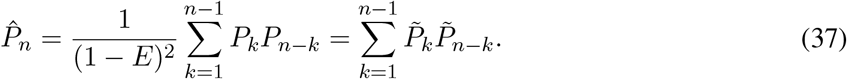

Since the 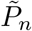 are identical between congruent scenarios (for any *n ≥* 1), we conclude that 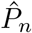 are also identical between congruent scenarios (for any *n ≥* 2).

#### S.1.5 Constructing congruent scenarios

For any given BDS scenario (*λ, µ, ψ*) there are various approaches towards constructing alternative congruent scenarios, described below. All of the presented methods have in common that one first calculates the “pulled variables” of the original scenario (which fully define the scenario’s congruence class), and then provides sufficient additional constraints to obtain a new single scenario within the congruence class. The discussion below also reveals how scenarios can become fully identifiable when additional constraints are provided.

##### S.1.5.1 Congruent scenarios by specifying ψ

In the first approach, we specify a new sampling rate *ψ*\* and then adjust the remaining model variables to obtain a congruent scenario. Specifically, let 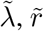, and 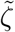 be the pulled variables of the scenario as defined in Supplement S.1.2. Let *ψ* >* 0 be some arbitrary alternative sampling rate. Define 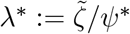 and:

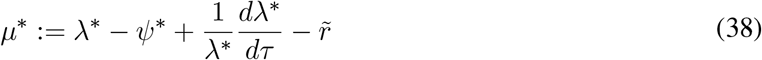

Then the pulled diversification rate of the new scenario (*λ*, µ*, ψ\**) is:

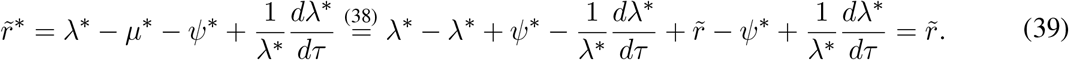

Further, 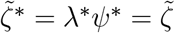. Hence, the new scenario has the same 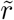 and the same 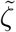 as the original scenario. By Supplement S.1.3, the two scenarios are thus congruent. Hence, simply by choosing an alternative sampling rate *ψ*, one can find a corresponding congruent scenario (*λ*, µ*, ψ\**), as long as the resulting *µ*\* is physically meaningful (i.e., non-negative). Reciprocally, if the congruence class of a diversification/epidemiological scenario is known (e.g., estimated via maximum-likelihood), and in addition *ψ* is somehow independently estimated, then the full scenario can be reconstructed.

### S.1.5.2 Congruent scenarios by specifying *µ* and *λ*(*τ*_*o*_)

Instead of first specifying an alternative *ψ*\*, one could construct congruent scenarios by first specifying *µ*\* and the speciation rate *λ*\*(*τ*_*o*_) at some age *τ*_*o*_, as follows. Let 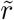 and 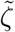 be the pulled variables of the scenario (*λ, µ, ψ*) as defined in Supplement S.1.2. For any given extinction rate profile *µ\**, age *τ*_*o*_ and speciation rate 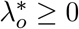 at age *τ*_*o*_, choose *λ*\* as the solution of the following differential equation:

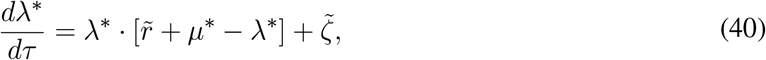

with condition 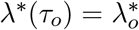, and set 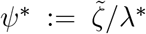 The pulled diversification rate of the new scenario (*λ*, µ*, ψ\**) is given by:

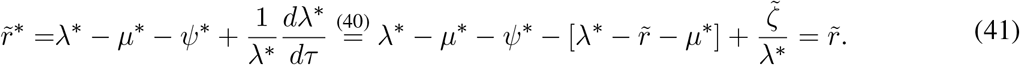

Further, 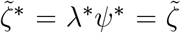. Hence, the new scenario has the same 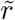 and the same 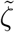 as the original scenario. By Supplement S.1.3, the two scenarios are thus congruent. Hence, simply by choosing an alternative sampling rate *µ*\* and an arbitrary initial condition *λ*\*(*τ*_*o*_) at some age *τ*_*o*_, one can find a corresponding congruent scenario (*λ*, µ*, ψ\**), provided that the resulting *λ* is physically meaningful (i.e., non-negative). Reciprocally, if the congruence class of a diversification/epidemiological scenario is known (e.g., estimated via maximum-likelihood), and in addition *µ* and *λ*(*τ*_*o*_) are somehow independently obtained (e.g., from clinical experiments), then the full scenario can be reconstructed.

Note that, as long as *λψ >* 0 and *λ*\*(0) *>* 0, and as long as 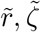 and *µ*\* are continuous, the corresponding *λ*\* will always be strictly positive. To see why this is the case, observe that if *λ*\* was zero or negative at some point, there would exist an age *τ*_z_ where *λ*\*(*τ*_z_) = 0 and *λ*\*(*τ*) *>* 0 for all *τ < τ*_z_. Since 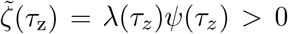 and 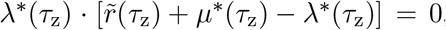, by Eq. (40) the derivative *dλ/dτ* must be strictly positive at *τ*_z_, and in fact strictly positive in a small neighborhood of *τ*_z_, which would mean *λ* could not reach zero at *τ*_z_ — a contradiction.

#### S.1.5.3 Congruent scenarios by specifying *µ* and *ψ*(*τ*_*o*_)

Alternatively to specifying the profile *µ*\* and the value *λ*\*(*τ*\*_*o*_) at some age *τ*_*o*_, one could also specify the profile *µ*\* and the value *ψ*\*(*τ*_*o*_). In that case, simply choose 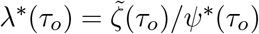, and apply the procedure in Supplement S.1.5.2 to obtain a congruent scenario (*λ*, µ*, ψ\**).

#### S.1.5.4 Congruent scenarios by specifying *λ*

Another way of constructing congruent scenarios is to first specify *λ* and then choose *µ* and *ψ* accordingly, as follows. Let 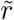 and 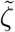 be the pulled variables of the scenario (*λ, µ, ψ*) as defined in Supplement S.1.2. For any given *λ*, choose 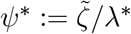 and:

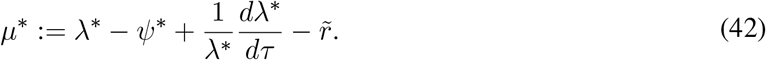

Then the pulled diversification rate of the new scenario (*λ*, µ*, ψ\**) is:

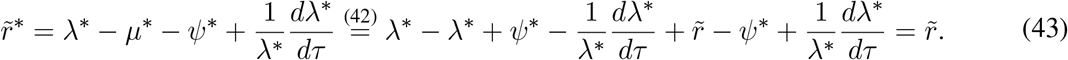

Hence, the new scenario has the same 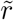 and the same 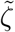 as the original scenario. The two scenarios are thus congruent. Hence, simply by choosing an alternative sampling rate *λ*, one can find a corresponding congruent scenario (*λ*, µ*, ψ\**), as long as the resulting *µ*\* is physically meaningful (i.e., non-negative).

#### S.1.5.5 Congruent scenarios by specifying *R*_e_ and *λ*(*τ*_*o*_)

Congruent scenarios can also be constructed by first choosing the effective reproduction ratio profile and the speciation rate *λ*\*(*τ*_*o*_) at some age *τ*_*o*_, as follows. Let 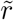 and 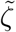 be the pulled variables of the scenario (*λ*, µ*, ψ\**) as defined in Supplement S.1.2. For any given basic reproduction ratio profile 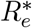, age *τ*_*o*_ and speciation rate 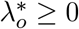 at age *τ*_*o*_, choose *λ* as the solution of the following differential equation:

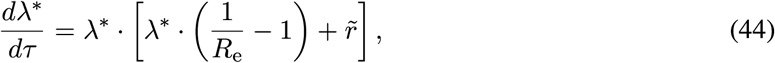

with condition 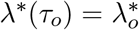, and set 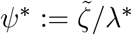 and *µ*\* :=λ*/*R*_e_ − *ψ*\*. The pulled diversification rate of the new scenario (*λ*, µ*, ψ\**) is given by:

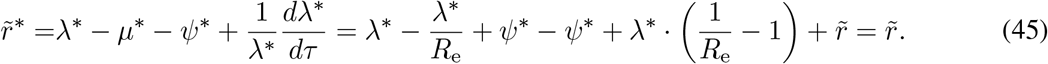

Further, 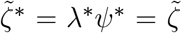. Hence, the new scenario has the same 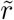 and the same 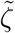 as the original scenario. By Supplement S.1.3, the two scenarios are thus congruent. Hence, simply by choosing an alternative 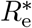 and an arbitrary condition 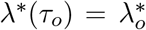 at some age *τ*_*o*_, one can find a corresponding congruent scenario (*λ*, µ*, ψ\**), provided that the resulting *λ* and *µ* are physically meaningful (i.e., non-negative). Reciprocally, if the congruence class of a diversification/epidemiological scenario is known (e.g., estimated via maximum-likelihood), and in addition *R*_e_ and *λ*\*(*τ*_o_) are somehow independently obtained, then the full scenario can be reconstructed.

Note that the differential equation (44) is of Bernoulli type, and can thus be solved analytically:

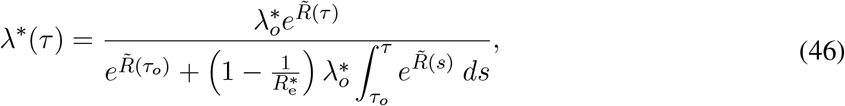

where 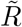 is defined as in Eq. (29).

#### S.1.5.6 Congruent scenarios by specifying *µ* + *ψ* and *λ*(*τ*_*o*_)

Congruent scenarios can also be constructed by first choosing the total removal rate *δ* := *µ* + *ψ* and the speciation rate *λ*(*τ*_*o*_) at some age *τ*_*o*_, as follows. Let 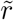 and 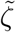 be the pulled variables of the scenario (*λ*, µ*, ψ\**) as defined in Supplement S.1.2. For any given profile *δ ≥* 0, age *τ*\*_*o*_ and speciation rate 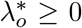 at age 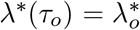, choose *λ*\* as the solution of the following differential equation:

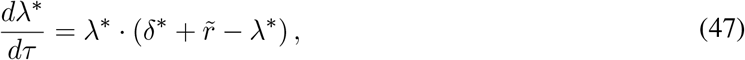

with condition 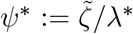, and set 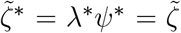 and 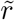. The pulled diversification rate of the new scenario (*λ*, µ*, ψ\**) is given by:

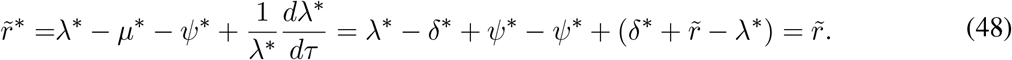

Further, 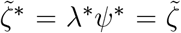. Hence, the new scenario has the same 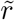 and the same 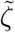 as the original scenario. By Supplement S.1.3, the two scenarios are thus congruent. Hence, simply by choosing an alternative *δ*\* and an arbitrary condition 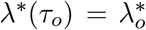 at some age *τ*_*o*_, one can find a corresponding congruent scenario (*λ*, µ*, ψ\**), provided that the resulting *λ*\* and *µ*\* are physically meaningful (i.e., non-negative). Reciprocally, if the congruence class of a diversification/epidemiological scenario is known (e.g., estimated via maximum-likelihood), and in addition *δ* and *λ*(*τ*_*o*_) are somehow independently obtained, then the full scenario can be reconstructed.

Note that the differential equation (47) is of Bernoulli type, and can thus be solved analytically:

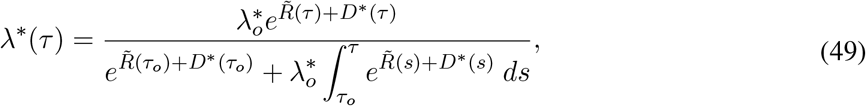

where 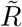 is defined as in Eq. (29) and *D* is defined as:

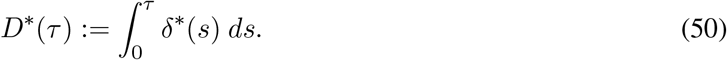

From Eq. (49) it becomes clear that if *τ*_o_ = 0 and 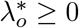, then *λ*\*(*τ*) will be non-negative for all *τ ≥* 0 (i.e., physically meaningful).

#### S 1.5.7 Congruent scenarios by specifying *µ* + *ψ* and *S*(*τ*_*o*_)

Congruent scenarios can also be constructed by first choosing the total removal rate *δ* := *µ* + *ψ* and the sampling proportion *S*(*τ*_*o*_) at some age *τ*_*o*_, as follows. Let 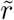 and 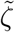 be the pulled variables of the scenario (*λ, µ, ψ*) as defined in Supplement S.1.2. For any given profile *δ >* 0, age *τ*_*o*_ and sampling proportion 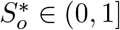 at age *τ*\*_*o*_, choose *λ*\* as the solution of the following differential equation:

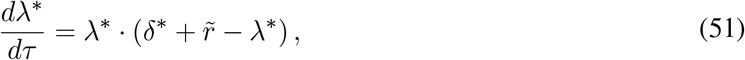

with initial condition 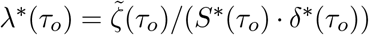, and set 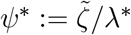 and *µ*\* := *δ*\* − *ψ*\*. The pulled diversification rate of the new scenario (*λ*, µ*, ψ\**) is given by:

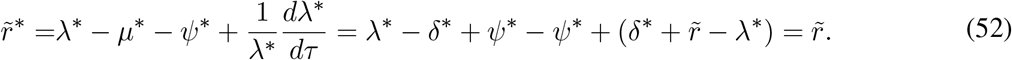

Further, 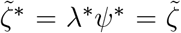. Hence, the new scenario has the same 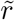 and the same 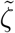 as the original scenario. By Supplement S.1.3, the two scenarios are thus congruent. Hence, simply by choosing an alternative *δ*and an arbitrary condition 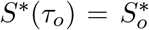 at some age *τ*_*o*_, one can find a corresponding congruent scenario (*λ*, µ*, ψ\**), provided that the resulting *λ*\* and *µ*\* are physically meaningful (i.e., non-negative). Reciprocally, if the congruence class of a diversification/epidemiological scenario is known (e.g., estimated via maximum-likelihood), and in addition *δ* and *S*(*τ*_*o*_) are somehow independently obtained, then the full scenario can be reconstructed.

#### S.1.5.8 Congruent scenarios by specifying *ψ/*(*µ* + *ψ*) and *λ*(*τ*_*o*_)

Congruent scenarios can also be constructed by first choosing the “sampling proportion” *S* := *ψ/*(*µ* + *ψ*) and the speciation rate *λ*(*τ*_*o*_) at some age *τ*_*o*_, as follows. Let 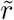 and 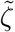 be the pulled variables of the scenario (*λ, µ, ψ*) as defined in Supplement S.1.2. For any given profile 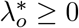, age *τ*_*o*_ and speciation rate *λ*\*_*o*_ 0 at age *τ*_*o*_, choose *λ* as the solution of the following differential equation:

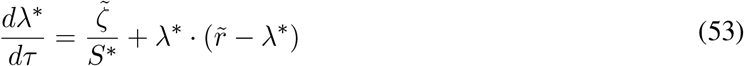

with condition 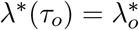, and set 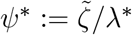 and *µ*\* := *ψ*/S\**− *ψ*\*. The pulled diversification rate of the new scenario (*λ*, µ*, ψ\**) is given by:

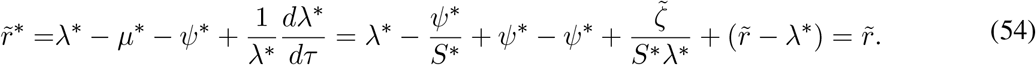

Further, 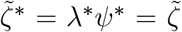. Hence, the new scenario has the same 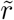 and the same 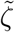 as the original scenario. By Supplement S.1.3, the two scenarios are thus congruent. Hence, simply by choosing an alternative *S* and an arbitrary condition 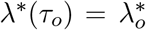 at some age *τ*_*o*_, one can find a corresponding congruent scenario (*λ*, µ*, ψ\**), provided that the resulting *λ* and *µ* are physically meaningful (i.e., non-negative). Reciprocally, if the congruence class of a diversification/epidemiological scenario is known (e.g., estimated via maximum-likelihood), and in addition *S* and *λ*(*τ*_*o*_) are somehow independently obtained, then the full scenario can be reconstructed.

#### S.1.6 The information content of concentrated sampling attempts

In the following we describe how samples obtained during concentrated sampling attempts can yield valuable insight into an epidemic. We define a concentrated sampling attempt (CSA) as a short period of sampling, taking place between two ages *τ*_1_ *> τ*_2_ *>* 0, and satisfying the following assumptions:

A. |*τ*_1_ − *τ*_2_| is much smaller than 1*/λ* and 1*/µ* throughout the CSA.
B. *λ* does not vary substantially between the times *τ*_1_ and *τ*_2_, i.e., we assume that *λ*(*τ*_1_) ≈ *λ*(*τ*_2_).
C. The probability that a lineage alive at *τ*_1_ will be sampled before age *τ*_2_, denoted *ρ*, is sufficiently high so that during the CSA the number of lineages sampled is much larger than the number of speciation and extinction events, i.e. *ρ* » *λ ·* |*τ*_1_ − *τ*_2_| and *ρ* » *µ ·* |*τ*_1_ − *τ*_2_|.
D. In the exterior vicinity of the open interval 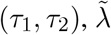 and 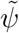 are continuous, so that 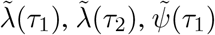 and 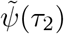 can be asymptotically identified.

We will show that in the limit of infinitely large trees, *λ*(*τ*_1_), *ψ*(*τ*_1_) and *ρ* can be accurately identified. Assuming that the CSA is so short that speciation and extinction are unlikely to occur in any given lineage (assumption A), we have:

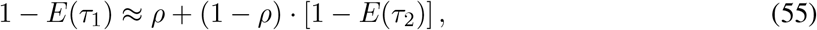

which is equivalent to:

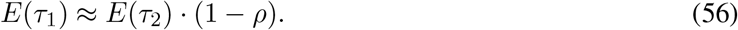

In other words, the probability that a lineage alive at age *τ*_1_ will eventually be sampled is approximately equal to the probability of being sampled during the CSA plus the probability of not being sampled during CSA multiplied by the probability that a lineage alive at age *τ*_2_ will eventually be sampled. In the absence of speciations and extinctions, the above approximation formula would be exact. Assuming that *λ*(*τ*_1_) ≈ *λ*(*τ*_2_) (assumption B) and using Eq. (56), we have:

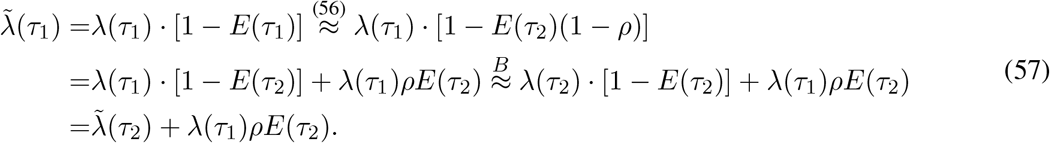

We mention that, strictly speaking, we only need to assume that *λ*(*τ*_1_) ≈ *λ*(*τ*_2_), i.e. in principle *λ* could vary during the CSA as long as it returns to its immediate pre-CSA value. The number of lineages sampled during the age interval [*τ*_1_, *τ*_2_], denoted *S*, can be accurately read off the tree, if the tree is sufficiently large. Assuming that speciations and extinctions within any given lineage are negligible compared to the sampling effort (i.e., changes in the total number of extant lineages are mostly due to sampling, assumption C), and for sufficiently large trees, we have:

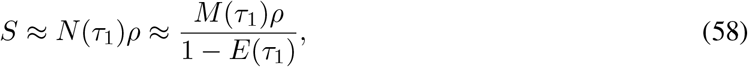

where *N* is the total number of extant lineages and *M* is the tree’s LTT. Denoting:

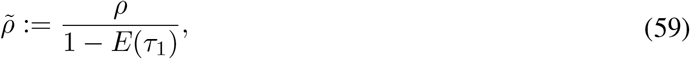

we see that 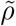 is asymptotically identifiable, since 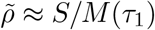. Note that:

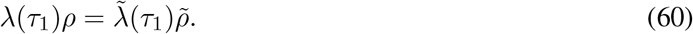

Using Eq. (57) and Eq. (60) we obtain:

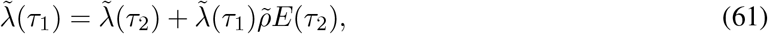

and hence:

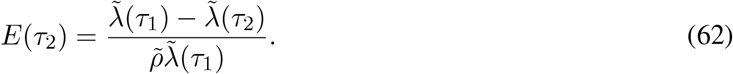

Since 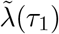 and 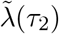 are asymptotically identifiable (assumption D), and 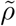 is also asymptotically identifiable, it follows that *E*(*τ*_2_) is also asymptotically identifiable. Note that the condition *τ*_2_ *>* 0 is necessary for ensuring that 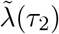 is asymptotically identifiable, since we need samples younger than *τ*_2_ for estimating 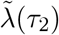. Solving Eq. (56) and Eq. (59) for *ρ* and *E*(*τ*_1_) leads to:

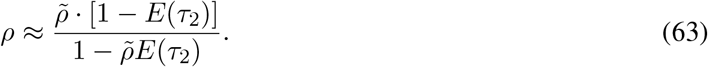

Hence, *ρ* is also asymptotically identifiable. Combined with Eq. (56), this implies that *E*(*τ*_1_) is also asymptotically identifiable. Solving Eq. (60) for *λ*(*τ*_1_) leads to:

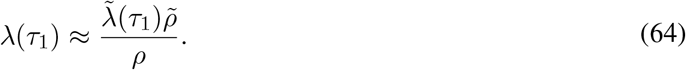

Hence, *λ*(*τ*_1_) is also asymptotically identifiable. Lastly, since 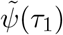 and 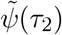 are asymptotically identifiable (assumption D), and since 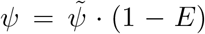, we conclude that *ψ*(*τ*_1_) and *ψ*(*τ*_2_) are also asymptotically identifiable.

#### S.1.7 Infinitesimally short concentrated sampling attempts

From a numerical perspective, it might be more practical to describe very short CSAs as separate instantaneous sampling processes with designated parameters (e.g., the time of the CSA and the probability of sampling a lineage during the CSA), rather than trying to approximate CSAs as very sharp peaks in *ψ*. The resulting model structure would then resemble the one described previously by Stadler *et al*. [10]. This is also the approach taken in the R package castor when fitting BDS models to timetrees: CSAs are parameterized separately from Poissonian sampling [44]. Formally, one can derive the corresponding likelihood starting from the BDS likelihood introduced earlier (Eqs. (3) or (6)) by considering a modified sampling rate of the form:

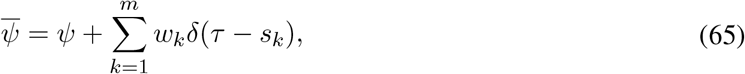

where *δ* is the Dirac distribution, *m* is the number of CSAs, *s*_*k*_ is the age of the *k*-th CSA (0 *≤s*_1_ *<* .. *< s*_*m*_ *≤ τ*_or_), *w*_*k*_ := − ln(1 − *ρ*_*k*_) is a weight, *ρ*_*k*_ *∈* (0, 1] is the sampling probability during the *k*-th CSA and ψ is the “background” (Poissonian) sampling rate outside of the CSAs. For a derivation of the corresponding likelihood of a bifurcating timetree in terms of *λ, µ, ψ*, the *s*_*k*_ and the *ρ*_*k*_ see [16]. For convenience, we provide the likelihood below using the notation of the present manuscript. We denote by *N*_*k*_ the number of tips sampled during and due to the *k*-th CSA, and by *n*_*k*_ the number of lineages “crossing” over *s*_*k*_ in the timetree. The likelihood density of the tree, when conditioned on the age of the origin *τ*_or_ and the survival of at least one lineage, is then given by:

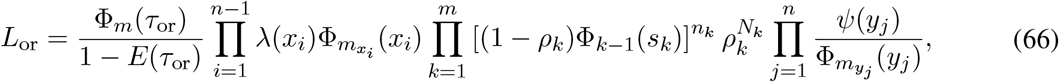

where as before *E*(*τ*) is the probability that a lineage alive at age *τ* is not included in the timetree, *m*_*τ*_ is the largest possible integer for which 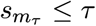, and Φ_0_, .., Φ_*m*_ are the sub-flows [58] between CSAs, defined as:

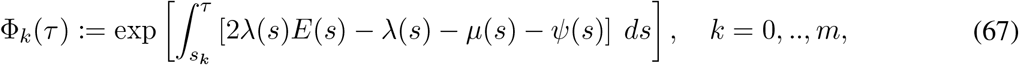

where for notational simplicity we defined *s*_0_ := 0. Note that for each *k* = 0, .., *m* the probability *E* satisfies within the age interval [*s*_*k*_, *s*_*k*+1_) the differential equation:

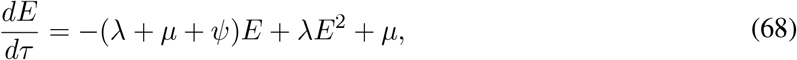

with initial condition 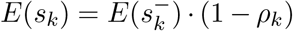 and *E*(0^−^) = 1. Similarly, one can derive the likelihood of a bifurcating timetree conditioned on the age of the tree’s root *τ*_r_, i.e., a splitting occurring at age *τ*_r_ and the eventual sampling of both child lineages:

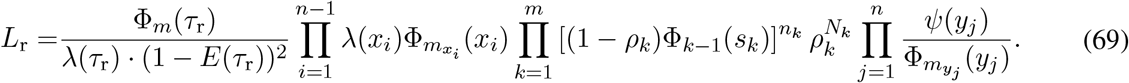

### S.2 Why conventional model selection methods cannot resolve congruencies

In this section we clarify why model selection methods such as AIC [45], BIC [59] or regularization [60], are not suitable for resolving model congruencies.

#### S.2.1 General considerations (big picture)

Model selection methods such as AIC, BIC and regularization essentially penalize excessive model complexity and are designed to prevent overfitting to finite datasets, i.e., to avoid inferring spurious complexities when the data at hand can also be adequately explained by simpler models. Indeed, it is common for multiple models to explain a dataset similarly well and in this case it is prudent to select the model with fewer free parameters since this tends to decrease the prediction error. These approaches do not, by any means, guarantee or even suggest that for any given dataset the selected (simpler) model is the one closest to the truth, as is often erroneously assumed. These methods merely constitute a methodological principle for gradually and conservatively choosing more complex models as new data appear, under the presumption that as new data accumulate the selected models will eventually converge to the truth.

When modelling pathogen epidemics, however, one must decide between congruent scenarios with differing levels of complexity, and no phylogenetic data, no matter how large, can distinguish between their proximity to the truth. In fact, all scenarios within a congruence class generate phylogenies with the same probability distribution, and hence they also have identical expected “prediction errors” (which, for example, AIC was designed to minimize). Hence, merely selecting the simpler model will almost always lead to an erroneous scenario, no matter how much sequence data we add. The very promise of conventional model selection — that selected models eventually converge to the truth as one adds more data — simply does not hold in this case.

One may of course compare scenarios belonging to different congruence classes since these will generally have different likelihoods for a particular dataset, and will certainly be distinguishable as the size of the dataset becomes very large. However, this does not solve the problem, since one is not really selecting the best model for the data but in fact merely the best congruence class. Even if one of the scenarios in the candidate set is entirely adequate for explaining the data at hand, or in the extreme case, is in the exact same congruence class as the true historical epidemiological dynamics, it does not follow that this scenario will resemble, even qualitatively, the true historical dynamics.

In conclusion, common model selection methods can at most be used to find a congruence class that “most efficiently” reproduces the data, or that has the lowest prediction error. In that sense, fitting congruence classes (e.g., by directly fitting pulled variables) is a parsimonious approach towards describing a phylogenetic dataset, while fitting full birth-death-sampling models adds redundant complexity that cannot possibly be identified using phylogenetic inference alone.

#### S.2.2 Regularization

Let us consider specifically the example of fitting piecewise linear (or piecewise constant) profiles for *λ, µ* and *ψ* on a grid over time, combined with a regularization approach that penalizes excessive oscillations or excessively large rate estimates (an example being Tikhonov regularization [61, 62]). To see why such an approach cannot possibly resolve model congruencies, suppose that a specific epidemiological history occurred, and consider the hypothetical limit where a phylogeny generated by that epidemic becomes infinitely large while the considered resolution of the fitted profiles (i.e., the number of grid points) remains fixed. In the absence of model congruencies, i.e., in typical inverse problems where regularization is commonly applied, one would expect that the fitted profiles would eventually approach the true historical profiles, provided of course that the grid resolution is high enough to approximately capture the true historical profiles. But this is clearly not the case here. Indeed, in the limit of an infinitely large phylogeny regularization becomes irrelevant, since for any given regularization parameter (also known as “smoothing parameter”) minimizing deviations from the data becomes infinitely more important than reducing the regularization penalties; in other words, for very large trees regularized profile fitting becomes equivalent to maximum-likelihood fitting. As we have demonstrated using simulations (Supplemental Figs. S1 and S2), maximum-likelihood-fitted profiles can be completely wrong even when using massive phylogenies with tens of thousands of tips and for relatively simple epidemiological histories. The situation can only be worse for smaller datasets.

#### S.2.3 AIC and BIC

Alternatively, let us consider the example of fitting various functional forms for *λ, µ* and *ψ* via maximum-likelihood, with the “best” functional form being selected using AIC [45] or BIC [59]. For any given choice of functional forms, the maximum-likelihood-fitted model will a priori tend to be the one closest to the congruence class of the true epidemiological history, rather than the true epidemiological history itself. Choosing the functional forms that minimize AIC or BIC would only yield a model that balances the number of parameters against the goodness of fit to the congruence class, but not against the goodness of fit to the true epidemiological history. There is little reason to expect that the fitted model selected via AIC or BIC will happen to actually be close to the true history, even for massive datasets (examples in Supplemental Figs. S1 and S2). The situation can only be worse for small datasets.

### S.3 Overview of computer code

This section provides an overview of computational tools developed as part of this manuscript, and publicly available in the R package `castor` v1.6.5 [44], for working with BDS models and congruencies. The functions described below have been designed with efficiency in mind, and can typically scale well to phylogenies with hundreds of thousands of tips. For detailed instructions consult the `castor` user manual.

#### generate_tree_hbds

Generate a random timetree according to a time-dependent BDS model with arbitrary rates *λ, µ, ψ* through time. The rates are specified as piecewise linear curves (or quadratic splines, or cubic splines) on a discrete time grid. By choosing a sufficiently fine time grid, any arbitrary functional forms can in principle be accommodated to arbitrary accuracy.

#### simulate_deterministic_hbds

Simulate a time-dependent BDS model with arbitrary rates *λ, µ, ψ* through time, in the deterministic limit (i.e., use differential equations rather than a stochastic process). This function can be used to calculate various alternative parameters of a BDS model, such as the *R*_e_, the removal rate *δ*, the sampling proportion *S*, the LTT, the pulled speciation rate 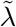 and so on. In particular, this function can be used to check if two BDS models are congruent, by comparing their pulled speciation rate 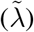 and event density (*λψ*).

#### fit_hbds_model_on_grid

Fit a time-dependent BDS model to a timetree via maximum-likelihood. The *λ, µ* and *ψ* can be assumed to be either piecewise constant, piecewise linear, quadratic splines or cubic splines defined on a discrete time grid; the values of these parameters are thus fitted at each grid point. This function uses parametric bootstrapping to calculate confidence intervals in the estimated parameters.

#### fit_hbds_model_parametric

Fit a BDS model to a timetree via maximum-likelihood. The profiles of *λ, µ* and *ψ* are described by user-specified functions that depend on a finite number of scalar parameters to be fitted. This function can thus be used to fit arbitrary functional forms. The function uses parametric bootstrapping to calculate confidence intervals in the estimated parameters.

#### congruent_hbds_model

Construct new BDS models by providing information on the congruence class (such as the pulled speciation rate) and any additional constraints needed for identifying a specific member of the congruence class. This function can be used to explore the set of models congruent to some reference model. It was used, for example, to generate the scenarios in Fig. 1 in the main article.

#### model_adequacy_hbds

Test if a given BDS model (or a distribution of BDS models) adequately describes a given timetree in terms of various statistics. For example, this function can perform a Kolmogorov-Smirnov test to examine if the distribution of node ages (or tip ages, or edge lengths) in the timetree deviate significantly from those expected under the model(s). Note that, due to the existence of model congruencies, even if a BDS model adequately describes a given timetree this does not mean that the model is even close to the original BDS scenario that generated the tree. If a BDS model adequately describes a timetree, it merely means that any of the myriad of members of the model’s congruence class would generate trees similar to the tree at hand.

---

**Figure S1:**
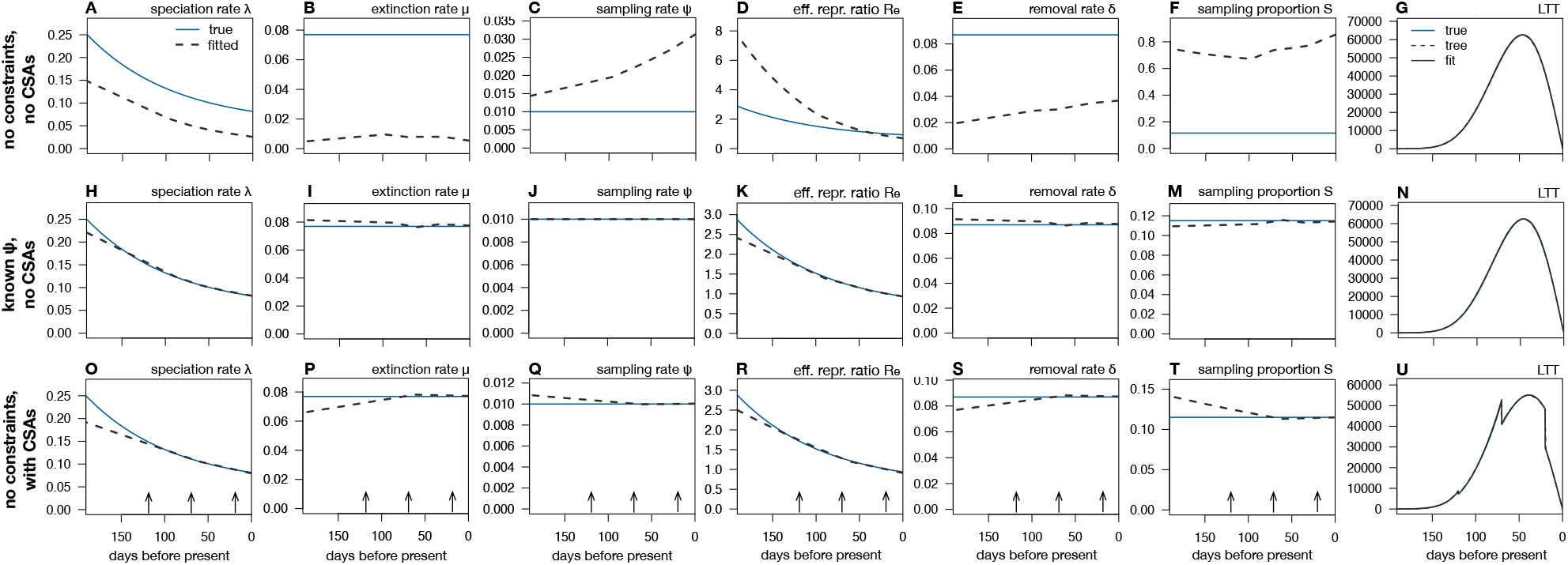
Limits to reconstructing an epidemic’s dynamics via maximum-likelihood (scenario 1). (A–F) Maximum-likelihood estimates (grey dashed curves) of the speciation rate (*λ*), extinction rate (*µ*), sampling rate (*ψ*), effective reproduction ratio (*R*_e_), removal rate (*δ* = *µ* + *ψ*) and sampling proportion (*S* = *ψ/*(*µ* + *ψ*)) over time, based on a timetree with 175,440 tips simulated under a hypothetical birth-death-sampling scenario (blue continuous curves) and without any additional constraints. All rates are in day^−1^. Model adequacy was confirmed using predictive posterior simulations with multiple tests. Observe the poor agreement between the estimated and true profiles. (G) Deterministic LTT (dLTT) of the fitted model, compared to the true scenario’s dLTT and the LTT of the timetree (note that all curves are nearly identical). (H–N) Similar to A–G, but for a model fitted to the same data as in A–G while fixing the sampling rate to its true profile. Observe the improved agreement between estimated and true profiles. (O–U) BDS parameters and dLTT for a model fitted to a timetree with 172,888 tips, generated under nearly the same BDS scenario as in A–G but with 3 added concentrated sampling attempts (CSAs, times indicated by vertical arrows). No additional constraints were used during fitting.

**Figure S2:**
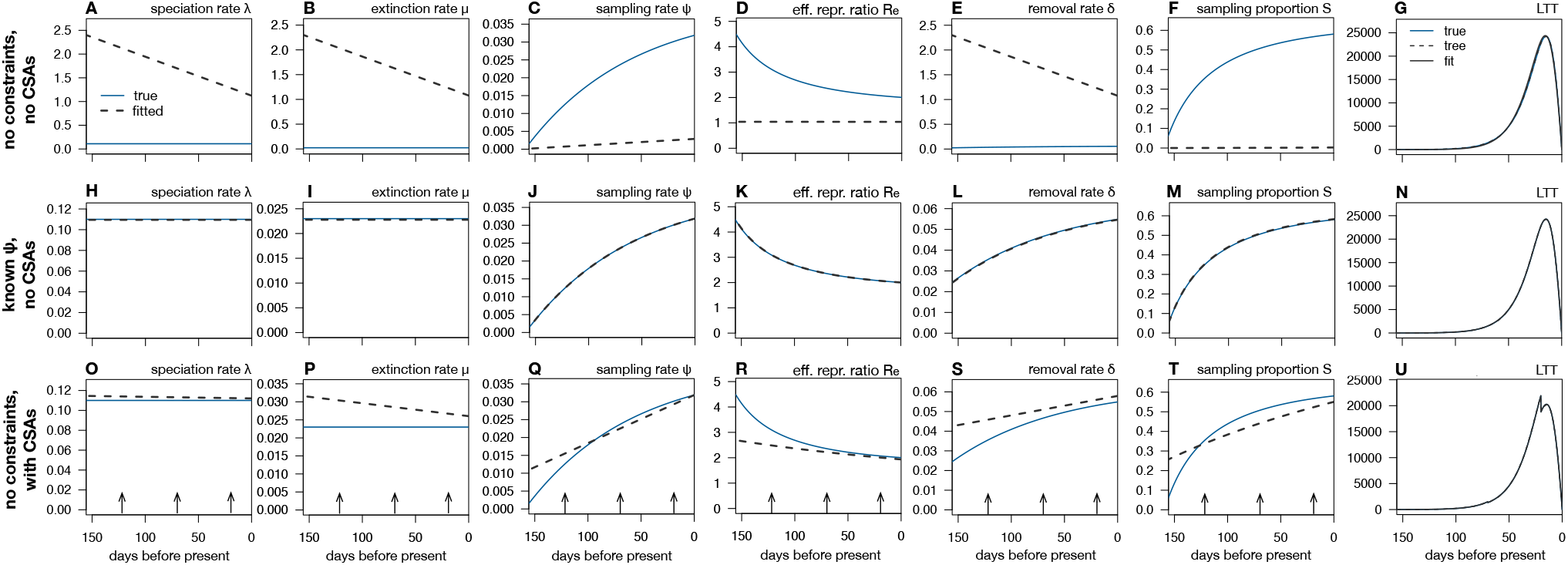
Limits to reconstructing an epidemic’s dynamics via maximum-likelihood (scenario 2). (A–F) Maximum-likelihood estimates (grey dashed curves) of the speciation rate (*λ*), extinction rate (*µ*), sampling rate (*ψ*), effective reproduction ratio (*R*_e_), removal rate (*δ* = *µ* + ψ) and sampling proportion (*S* = *ψ/*(*µ* + *ψ*)) over time, based on a timetree with 55,934 tips simulated under a hypothetical birth-death-sampling scenario (blue continuous curves) and without any additional constraints. All rates are in day^−1^. Model adequacy was confirmed using predictive posterior simulations with multiple tests. Observe the poor agreement between the estimated and true profiles. (G) Deterministic LTT (dLTT) of the fitted model, compared to the true scenario’s dLTT and the LTT of the timetree (note that all curves are nearly identical). (H–N) Similar to A–G, but for a model fitted to the same data as in A–G while fixing the sampling rate to its true profile. Observe the improved agreement between estimated and true profiles. (O–U) BDS parameters and dLTT for a model fitted to a timetree with 51,619 tips, generated under nearly the same BDS scenario as in A–G but with 3 added concentrated sampling attempts (CSAs, times indicated by vertical arrows). No additional constraints were used during fitting.

**Figure S3:**
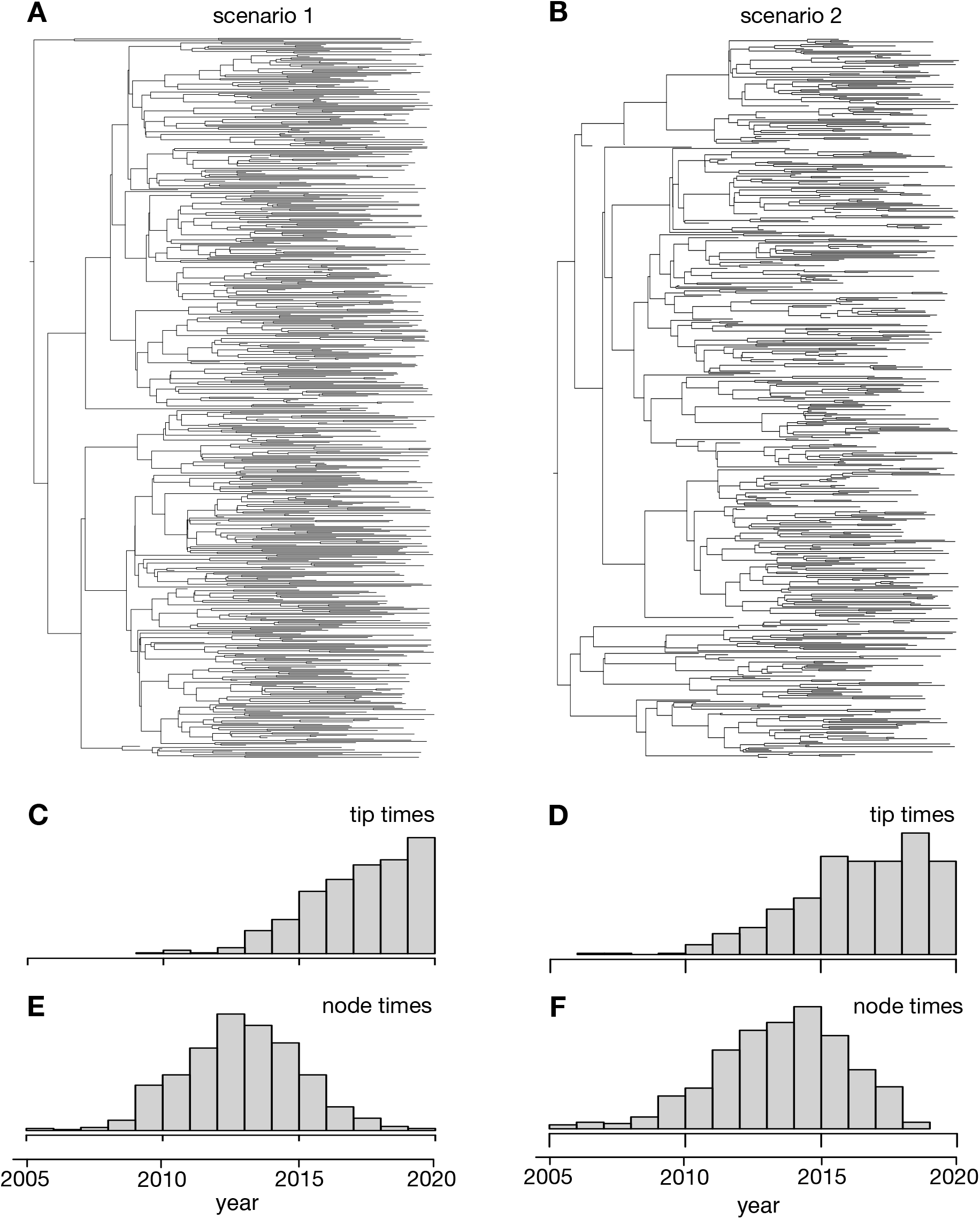
Phylogenetic trees simulated for the BEAST analysis. (A–B) Phylogenetic trees simulated under two hypothetical epidemiological scenarios, for Bayesian inference. (C, E) Histograms of tip ages and node ages in tree A (bar heights are in arbitrary units). (D, F) Histograms of tip ages and node ages in tree B (bar heights are in arbitrary units).

**Figure S4:**
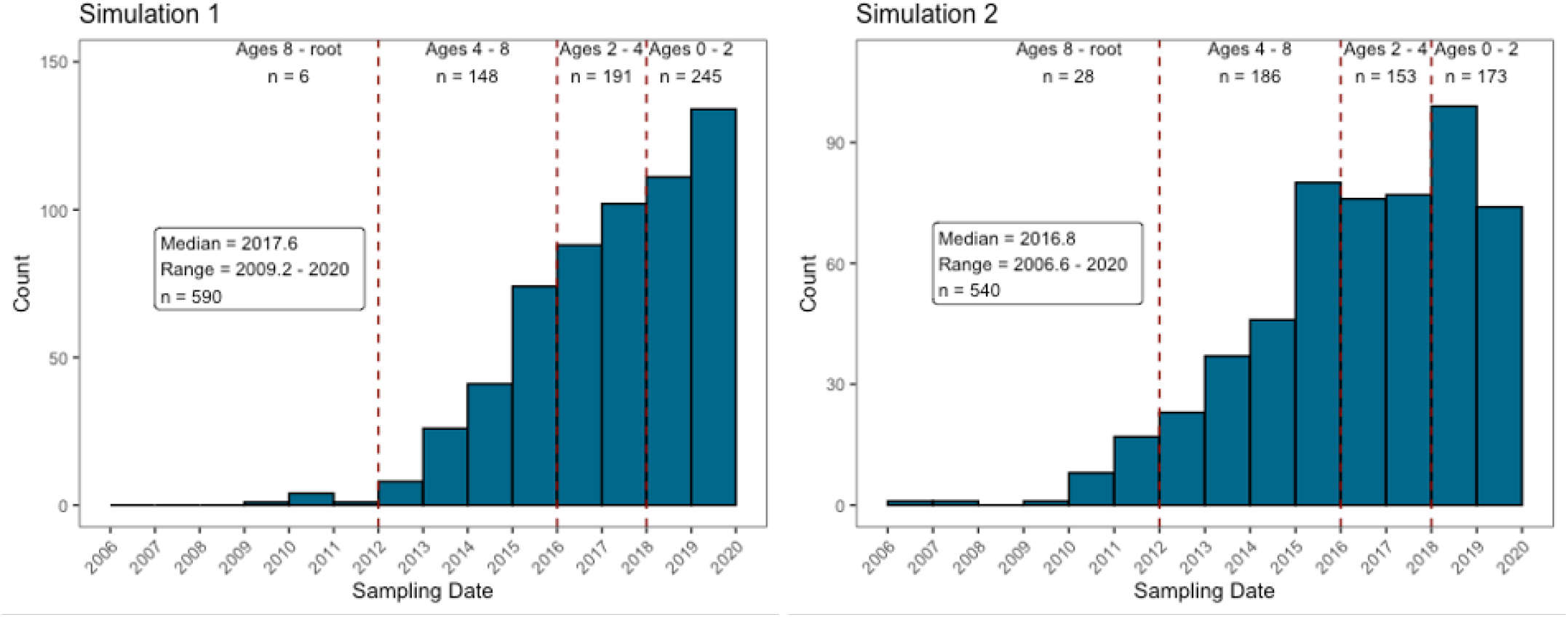
Choosing rate shift times for the BEAST simulation analysis. The distribution of sampling dates in the simulated trees informed the timing of specified rate shifts in fitted skyline models, to partially equalize the number of samples informing each interval and to ensure that the earliest interval had greater than zero samples.

**Figure S5:**
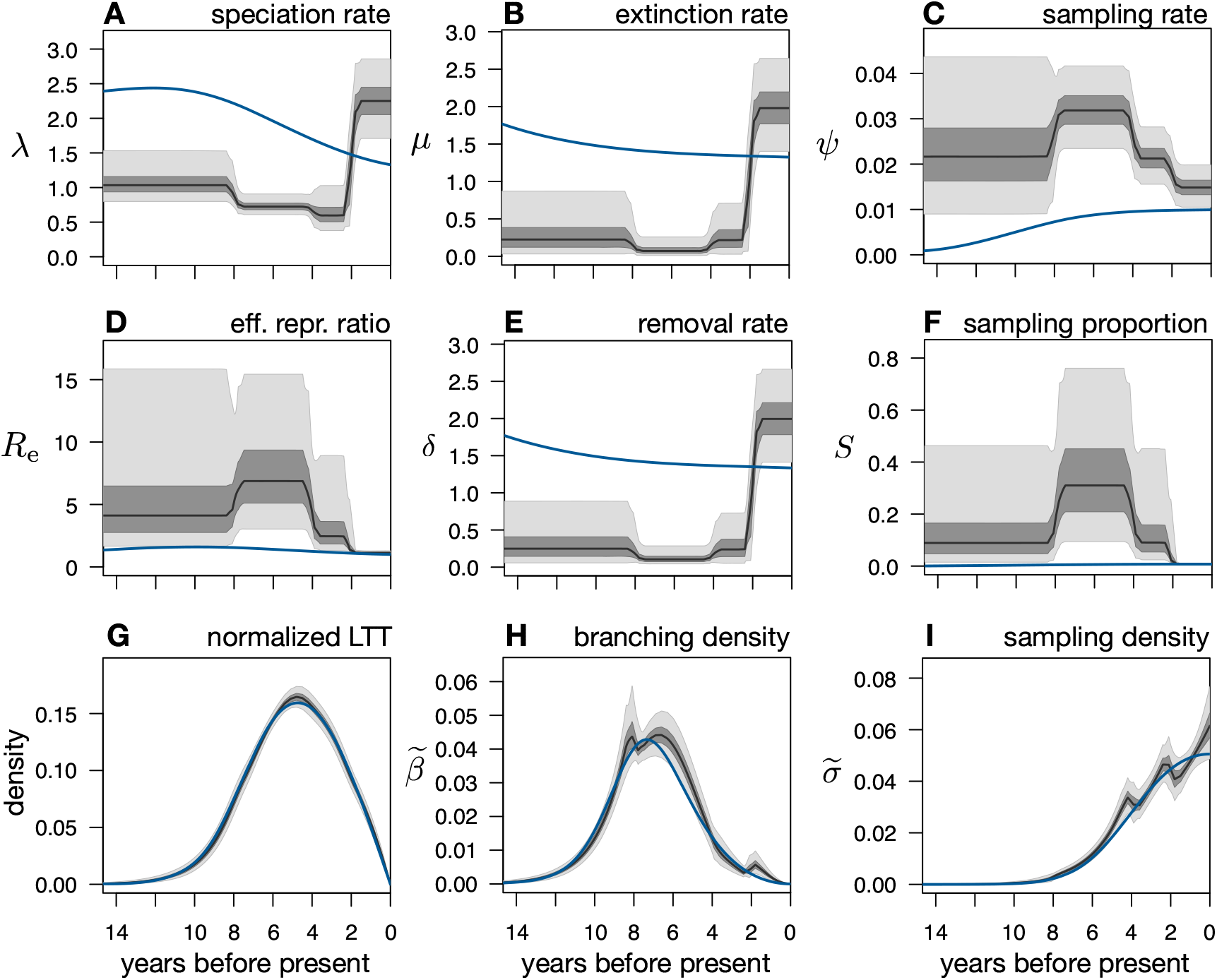
Reconstructing an epidemic’s dynamics in a Bayesian framework (BEAST2 run 1_444f). (A–F) Posterior distributions of the speciation rate (*λ*), extinction rate (*µ*), sampling rate (*ψ*), effective reproduction ratio (*R*_e_), removal rate (*δ* = *µ* + *ψ*) and sampling proportion (*S* = *ψ/*(*µ* + *ψ*)), as inferred from 590 sequences simulated under a hypothetical birth-death-sampling scenario (blue curves) using BEAST2 (tree in Supplemental Fig. S3A). Black curves show posterior median, dark and light shades represent equal-tailed 50%- and 95%-credible intervals of the posterior. All rates are in yr^−1^. The present-day sampling proportion was fixed to its true value during fitting, to account for previously reported identifiability issues [10]. Model adequacy was confirmed using predictive posterior simulations with multiple test statistics. Note the poor agreement between the predicted and true profiles. (G–I) Distributions of the deterministic lineages-through-time curves (normalized to unit area under the curve), branching densities 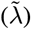 and sampling densities 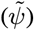, corresponding to the same posterior scenarios as in A–F, compared to their true profiled (blue curves). The relatively good agreement between the inferred and true profiles shows that BEAST2 closely reconstructed the epidemiological history’s congruence class but not the epidemiological history itself. See Supplemental Fig. S10 for the corresponding posterior distributions of the molecular evolution parameters.

**Figure S6:**
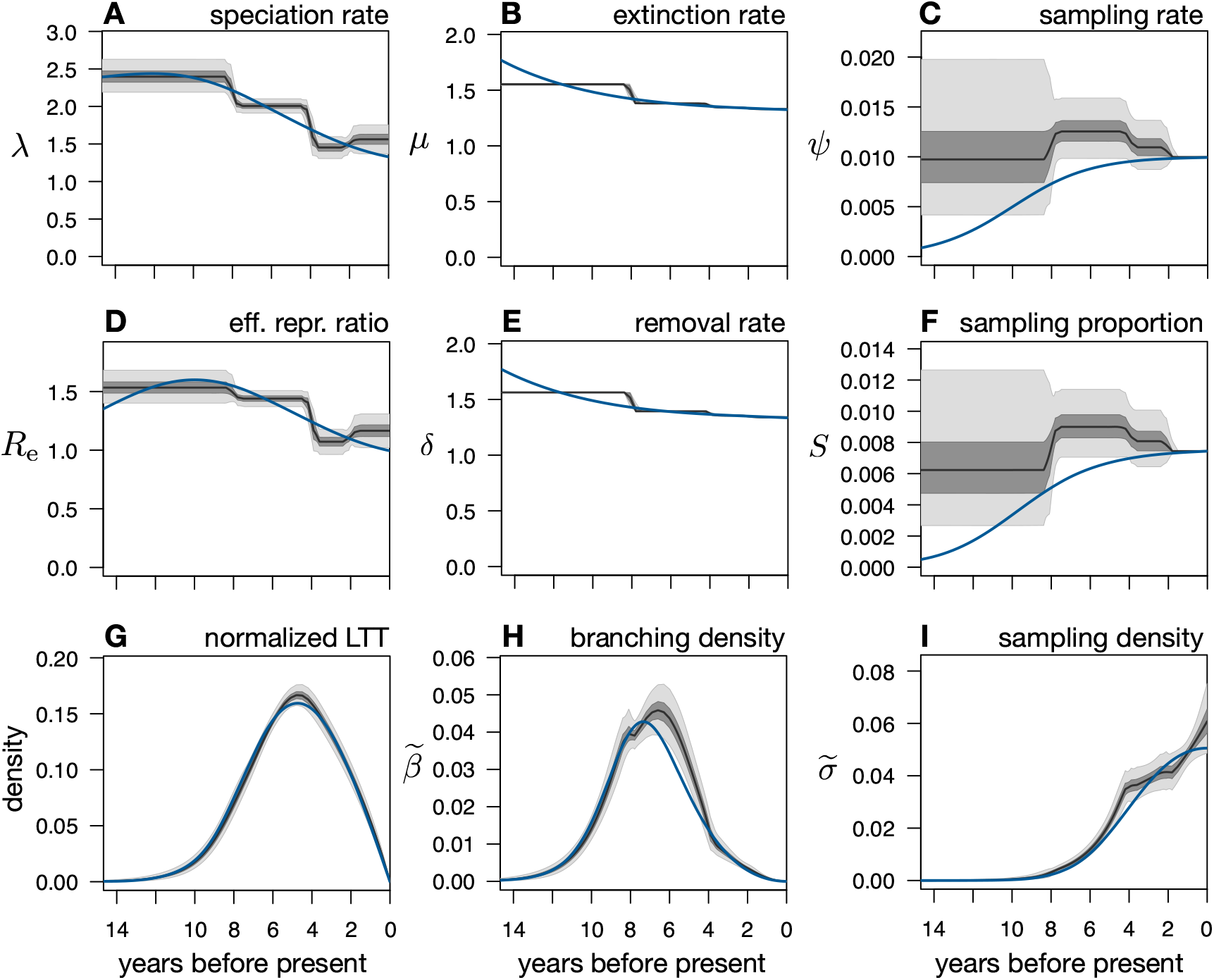
Reconstructing an epidemic’s dynamics in a Bayesian framework (BEAST2 run 1_44F4f). (A–F) Posterior distributions of the speciation rate (*λ*), extinction rate (*µ*), sampling rate (*ψ*), effective reproduction ratio (*R*_e_), removal rate (*δ* = *µ* + *ψ*) and sampling proportion (*S* = *ψ/*(*µ* + *ψ*)), as inferred from 590 sequences simulated under a hypothetical birth-death-sampling scenario (blue curves) using BEAST2 (tree in Supplemental Fig. S3A). Black curves show posterior median, dark and light shades represent equal-tailed 50%- and 95%-credible intervals of the posterior. All rates are in yr^−1^. The present-day sampling proportion as well as the removal rate (all intervals) were fixed to their true values during fitting. Model adequacy was confirmed using predictive posterior simulations with multiple test statistics. Note the much better agreement with the true profiles, compared to the situation where the removal rate was not fixed (Supplemental Fig. S5). (G–I) Distributions of the deterministic lineages-through-time curves (normalized to unit area under the curve), branching densities 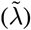 and sampling densities 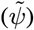, corresponding to the same posterior scenarios as in A–F, compared to their true profiled (blue curves). See Supplemental Fig. S11 for the corresponding posterior distributions of the molecular evolution parameters.

**Figure S7:**
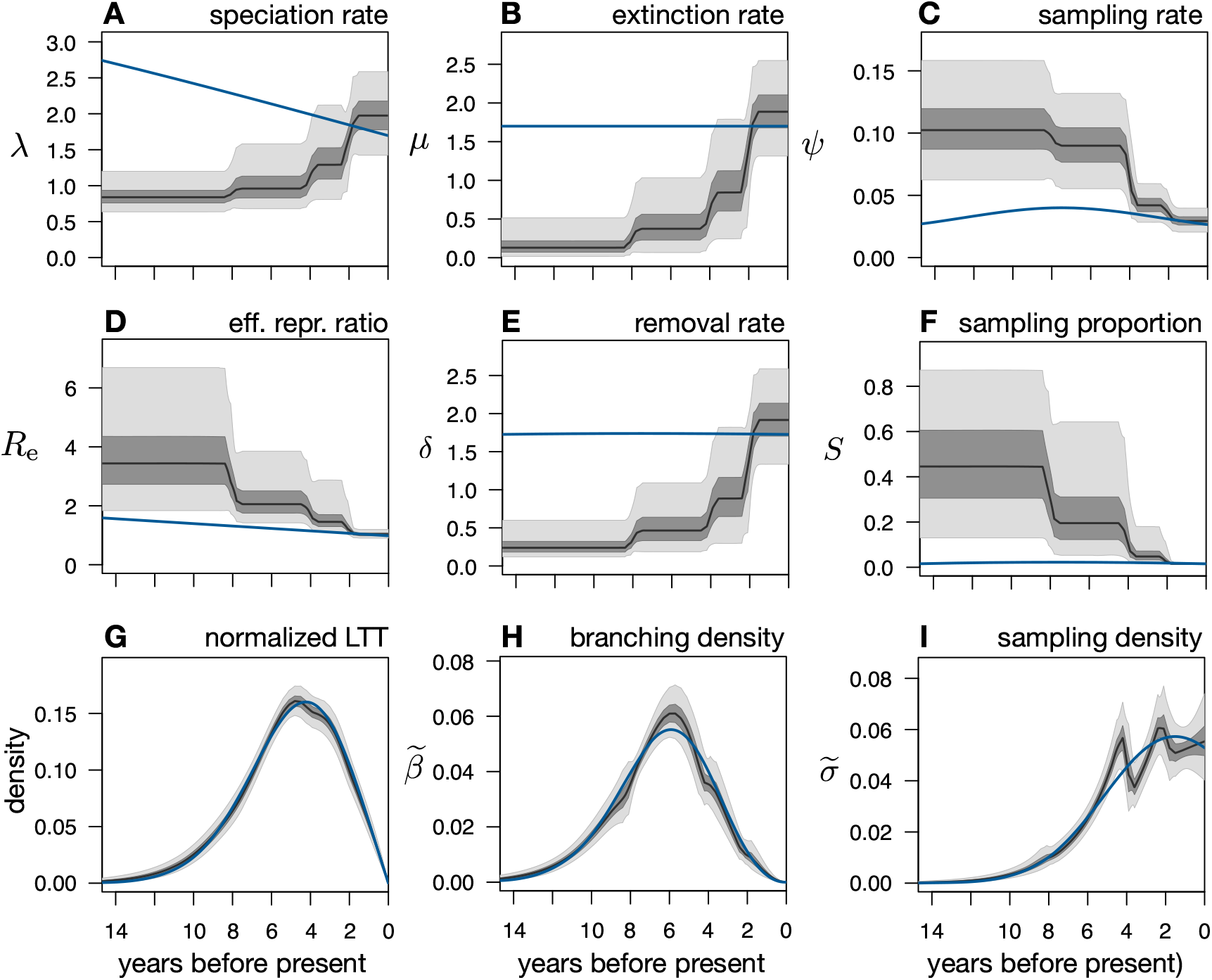
Reconstructing an epidemic’s dynamics in a Bayesian framework (BEAST2 run 2_444f). (A–F) Posterior distributions of the speciation rate (*λ*), extinction rate (*µ*), sampling rate (*ψ*), effective reproduction ratio (*R*_e_), removal rate (*δ* = *µ* + *ψ*) and sampling proportion (*S* = *ψ/*(*µ* + *ψ*)), as inferred from 540 sequences simulated under a hypothetical birth-death-sampling scenario (blue curves) using BEAST2 (tree in Supplemental Fig. S3B). Black curves show posterior median, dark and light shades represent equal-tailed 50%- and 95%-credible intervals of the posterior. All rates are in yr^−1^. The present-day sampling proportion was fixed to its true value during fitting, to account for previously reported identifiability issues [10]. Model adequacy was confirmed using predictive posterior simulations with multiple test statistics. Note the poor agreement between the predicted and true profiles. (G–I) Distributions of the deterministic lineages-through-time curves, branching densities 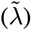 and sampling densities 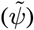, corresponding to the same posterior scenarios as in A–F, compared to their true profiled (blue curves). The relatively good agreement between the inferred and true profiles shows that BEAST2 closely reconstructed the epidemiological history’s congruence class but not the epidemiological history itself. See Supplemental Fig. S12 for the corresponding posterior distributions of the molecular evolution parameters.

**Figure S8:**
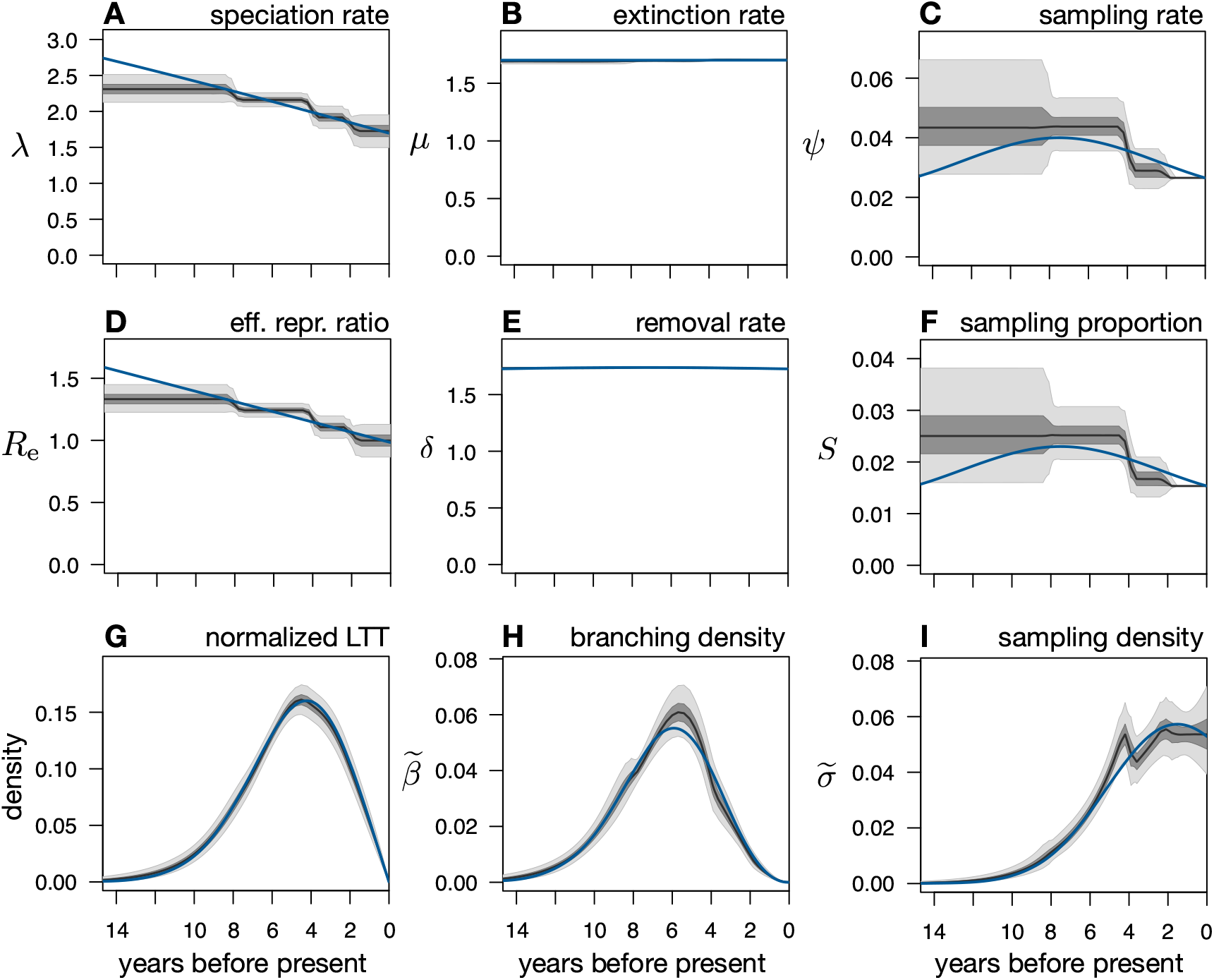
Reconstructing an epidemic’s dynamics in a Bayesian framework (BEAST2 run 2_44F4f). (A–F) Posterior distributions of the speciation rate (*λ*), extinction rate (*µ*), sampling rate (*ψ*), effective reproduction ratio (*R*_e_), removal rate (*δ* = *µ* + *ψ*) and sampling proportion (*S* = *ψ/*(*µ* + *ψ*)), as inferred from 540 sequences simulated under a hypothetical birth-death-sampling scenario (blue curves) using BEAST2 (tree in Supplemental Fig. S3B). Black curves show posterior median, dark and light shades represent equal-tailed 50%- and 95%-credible intervals of the posterior. All rates are in yr^−1^. The present-day sampling proportion and the removal rate (all time intervals) were fixed to their true value during fitting. Model adequacy was confirmed using predictive posterior simulations with multiple test statistics. Note the good agreement between the predicted and true profiles, made possible by fixing one of the model’s parameters. (G–I) Distributions of the deterministic lineages-through-time curves (normalized to unit area under the curve), branching densities 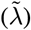 and sampling densities 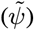, corresponding to the same posterior scenarios as in A–F, compared to their true profiled (blue curves). See Supplemental Fig. S14 for the corresponding posterior distributions of the molecular evolution parameters.

**Figure S9:**
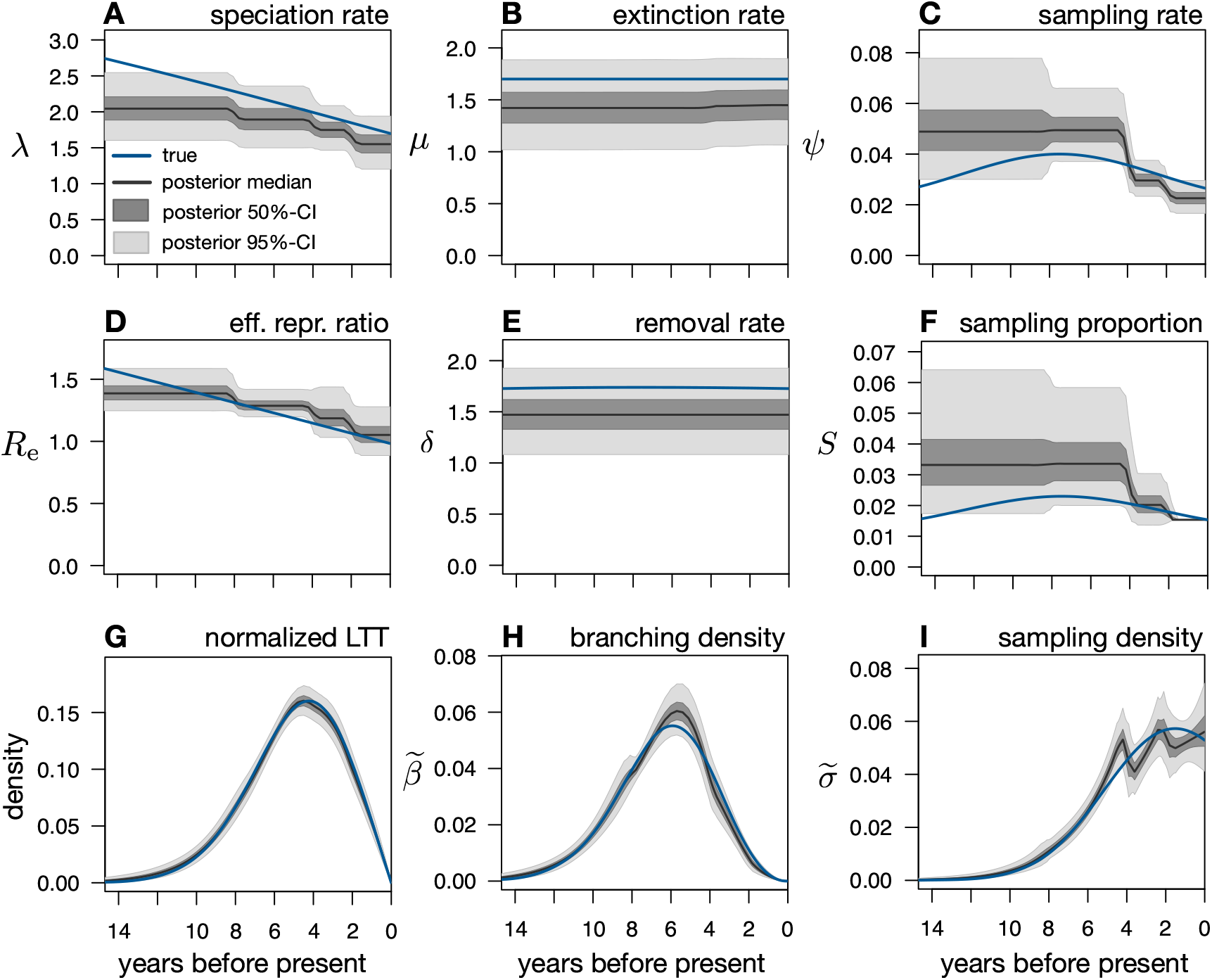
Reconstructing an epidemic’s dynamics in a Bayesian framework (BEAST2 run 2_414f). (A–F) Posterior distributions of the speciation rate (*λ*), extinction rate (*µ*), sampling rate (*ψ*), effective reproduction ratio (*R*_e_), removal rate (*δ* = *µ* + *ψ*) and sampling proportion (*S* = *ψ/*(*µ* + *ψ*)), as inferred from 540 sequences simulated under a hypothetical birth-death-sampling scenario (blue curves) using BEAST2 (tree in Supplemental Fig. S3B). Black curves show posterior median, dark and light shades represent equal-tailed 50%- and 95%-credible intervals of the posterior. All rates are in yr^−1^. The removal rate was constrained to be constant over time. The present-day sampling proportion was fixed to its true value during fitting, to account for previously reported identifiability issues [10]. Model adequacy was confirmed using predictive posterior simulations with multiple test statistics. (G–I) Distributions of the deterministic lineages-through-time curves (normalized to unit area under the curve), branching densities 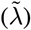 and sampling densities 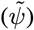, corresponding to the same posterior scenarios as in A–F, compared to their true profiled (blue curves). See Supplemental Fig. S13 for the corresponding posterior distributions of the molecular evolution parameters.

**Figure S10:**
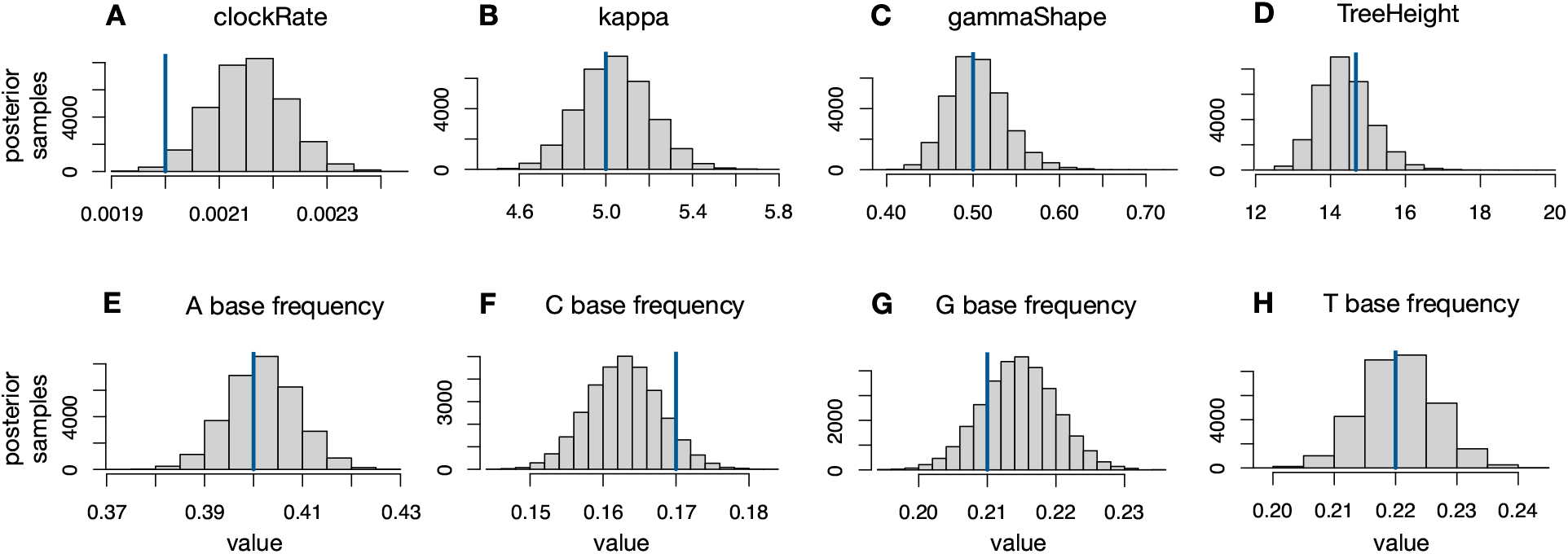
Posterior distributions of molecular evolution parameters (BEAST2 run 1_444f). Posterior distributions of (A) the mean molecular clock rate, (B) the transition/transversion ratio *κ*, (C) the shape parameter of the discretized gamma distribution of substitution rates, (D) the height of the tree or root age, and (E–H) the stationary frequencies of nucleotide bases A,C,G,T, as inferred from 590 sequences simulated under a hypothetical birth-death-sampling scenario using BEAST2 (epidemiological parameters in Supplemental Fig. S5, tree in Supplemental Fig. S3A). Histogram bars show frequencies. Blue vertical lines show the true values used in the simulation.

**Figure S11:**
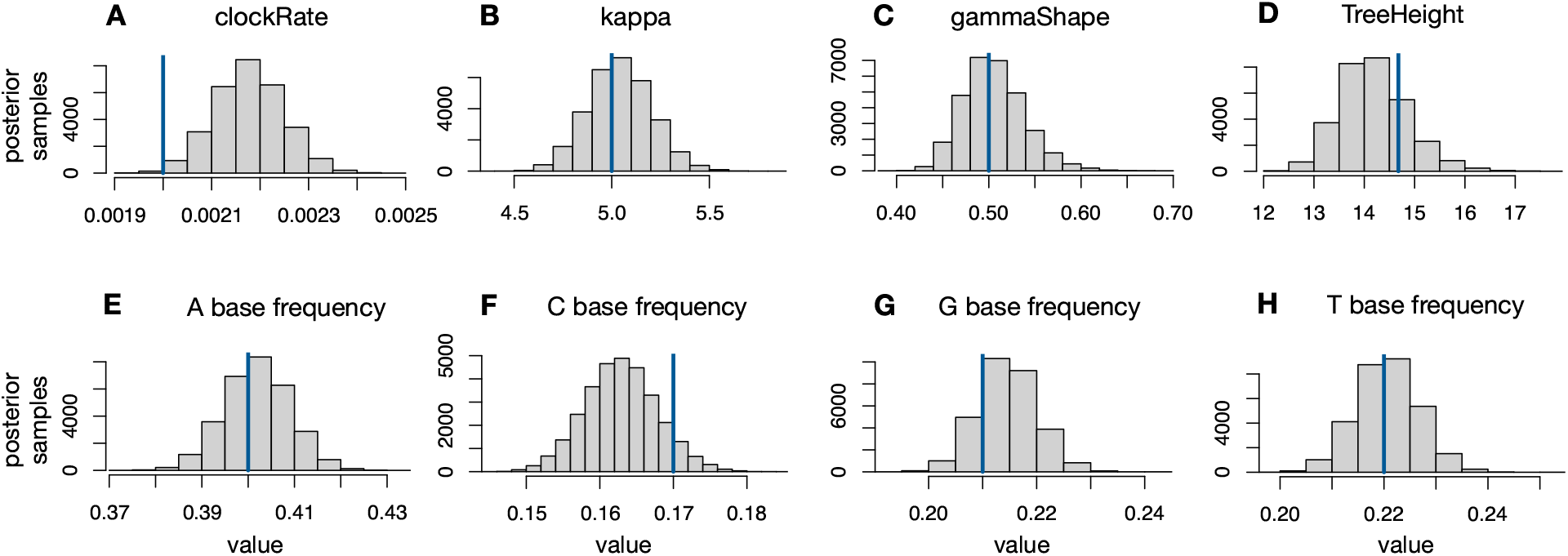
Posterior distributions of molecular evolution parameters (BEAST2 run 1_44F4f). Posterior distributions of (A) the mean molecular clock rate, (B) the transition/transversion ratio *κ*, (C) the shape parameter of the discretized gamma distribution of substitution rates, (D) the height of the tree or root age, and (E–H) the stationary frequencies of nucleotide bases A,C,G,T, as inferred from 590 sequences simulated under a hypothetical birth-death-sampling scenario using BEAST2 (epidemiological parameters in Supplemental Fig. S6, tree in Supplemental Fig. S3A). Histogram bars show frequencies. Blue vertical lines show the true values used in the simulation.

**Figure S12:**
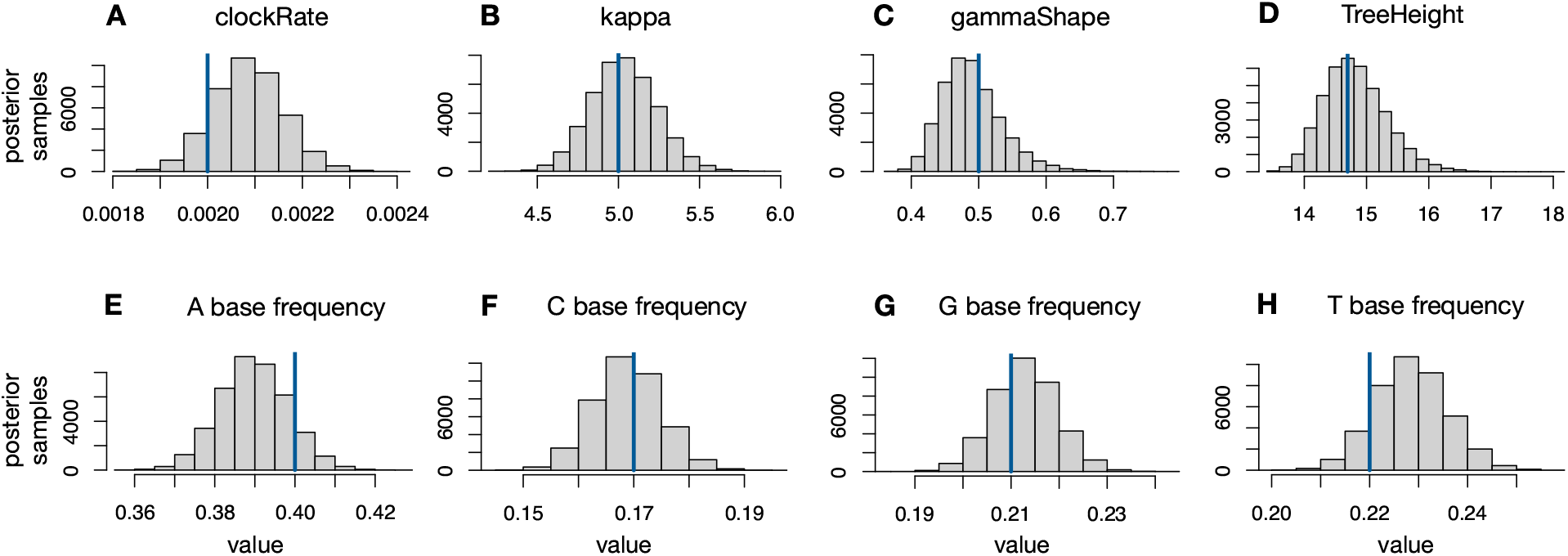
Posterior distributions of molecular evolution parameters (BEAST2 run 2_444f). Posterior distributions of (A) the mean molecular clock rate, (B) the transition/transversion ratio *κ*, (C) the shape parameter of the discretized gamma distribution of substitution rates, (D) the height of the tree or root age, and (E–H) the stationary frequencies of nucleotide bases A,C,G,T, as inferred from 540 sequences simulated under a hypothetical birth-death-sampling scenario using BEAST2 (epidemiological parameters in Supplemental Fig. S7, tree in Supplemental Fig. S3B). Histogram bars show frequencies. Blue vertical lines show the true values used in the simulation.

**Figure S13:**
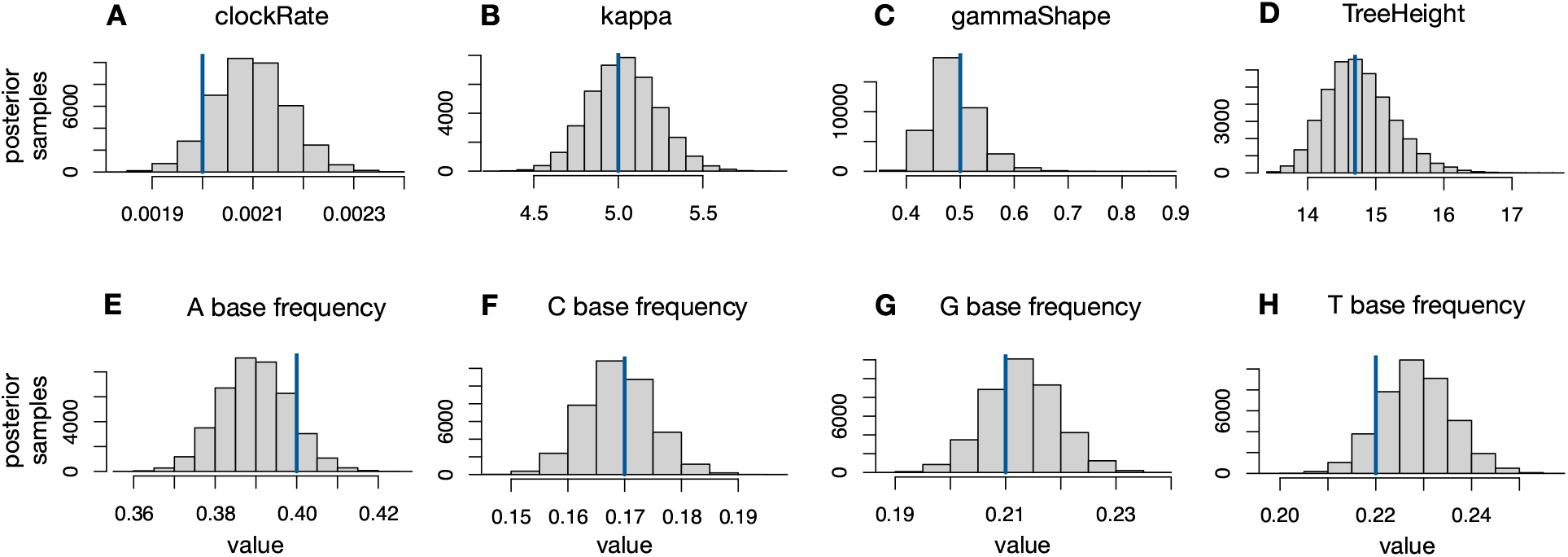
Posterior distributions of molecular evolution parameters (BEAST2 run 2_414f). Posterior distributions of (A) the mean molecular clock rate, (B) the transition/transversion ratio *κ*, (C) the shape parameter of the discretized gamma distribution of substitution rates, (D) the height of the tree or root age, and (E–H) the stationary frequencies of nucleotide bases A,C,G,T, as inferred from 540 sequences simulated under a hypothetical birth-death-sampling scenario using BEAST2 (epidemiological parameters in Supplemental Fig. S9, tree in Supplemental Fig. S3B). Histogram bars show frequencies. Blue vertical lines show the true values used in the simulation.

**Figure S14:**
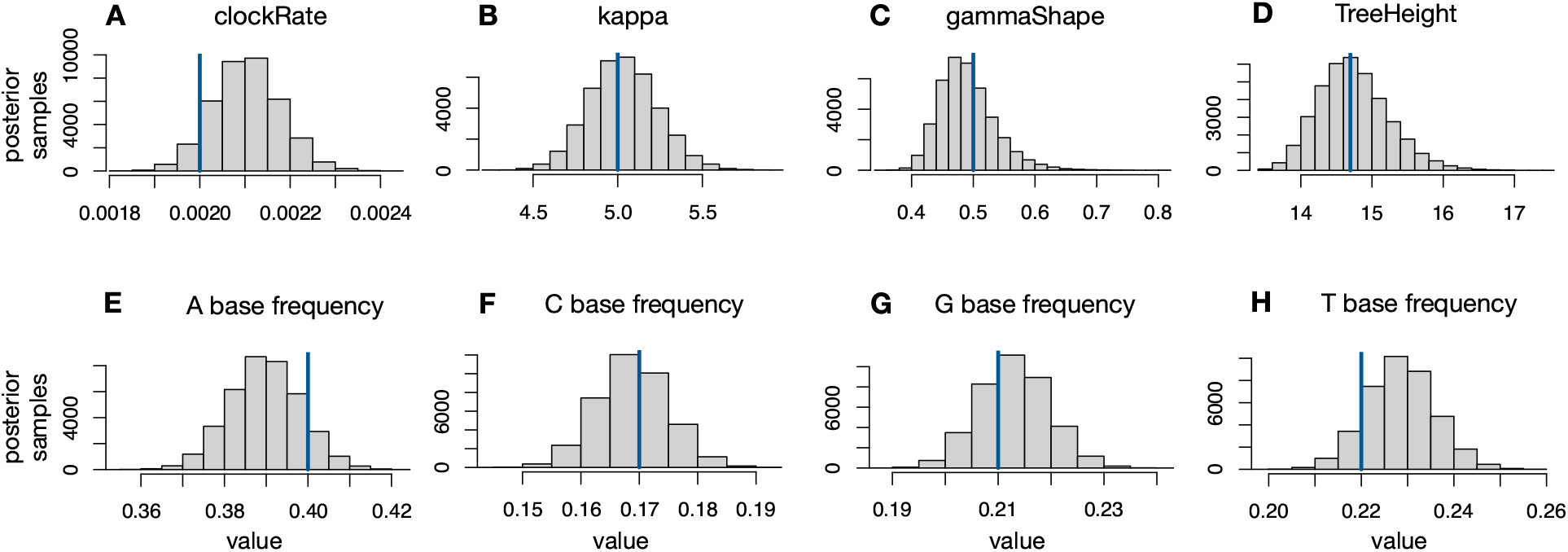
Posterior distributions of molecular evolution parameters (BEAST2 run 2_44F4f). Posterior distributions of (A) the mean molecular clock rate, (B) the transition/transversion ratio *κ*, (C) the shape parameter of the discretized gamma distribution of substitution rates, (D) the height of the tree or root age, and (E–H) the stationary frequencies of nucleotide bases A,C,G,T, as inferred from 540 sequences simulated under a hypothetical birth-death-sampling scenario using BEAST2 (epidemiological parameters in Supplemental Fig. S8, tree in Supplemental Fig. S3B). Histogram bars show frequencies. Blue vertical lines show the true values used in the simulation.

**Figure S15:**
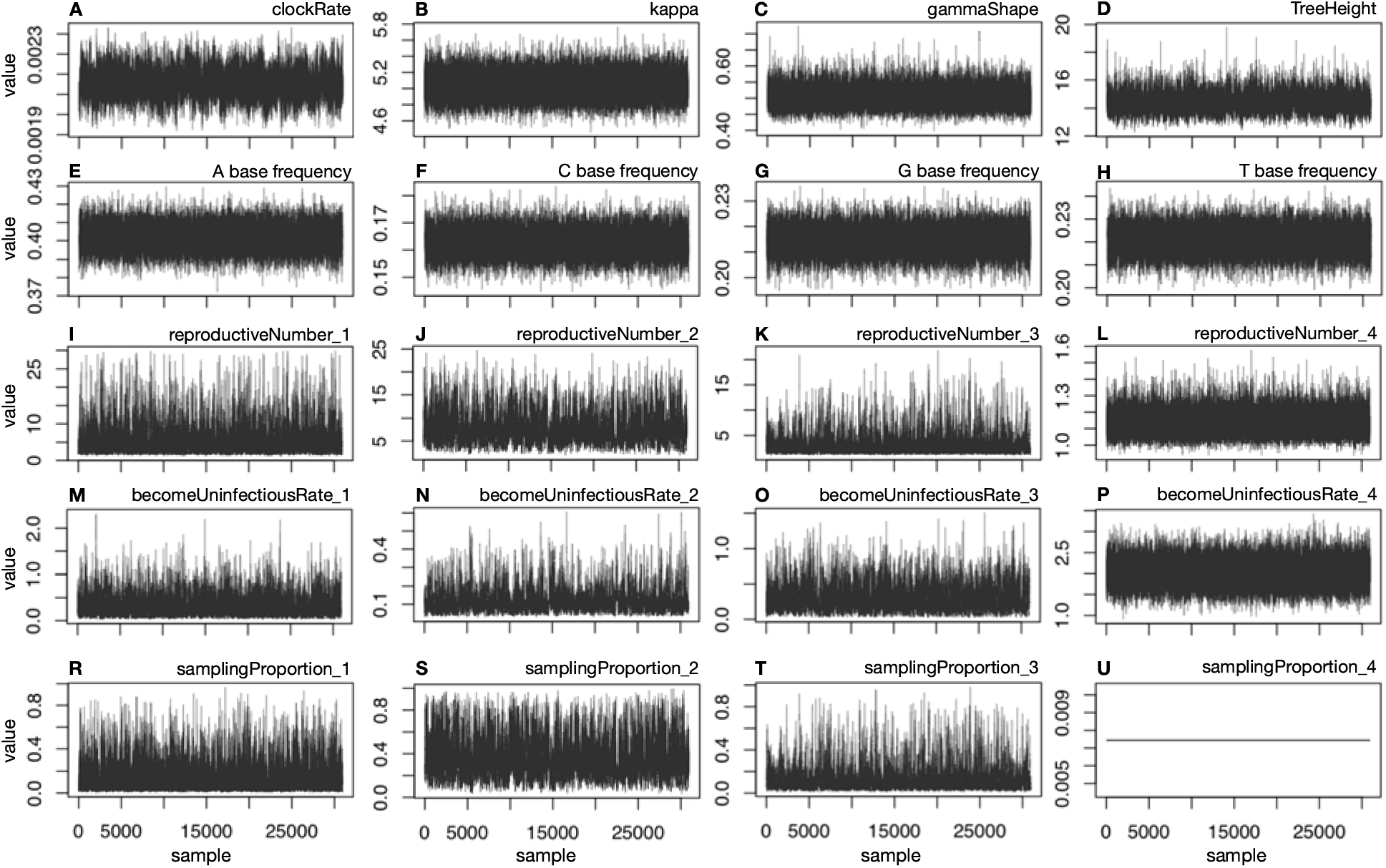
MCMC traces of molecular evolution and epidemiological parameters (BEAST2 run 1_444f). MCMC trace plots of molecular evolution and epidemiological (birth-death-sampling skyline model) parameters generated by BEAST2 (2 independent MCMC chains), based on the sequences simulated under the 1st hypothetical birth-death-sampling scenario (epidemiological parameters in Supplemental Fig. S5, tree in Supplemental Fig. S3A). Samples are shown after burn-in removal and after thinning of each MCMC chain, and after concatenating the two MCMC chains.

**Figure S16:**
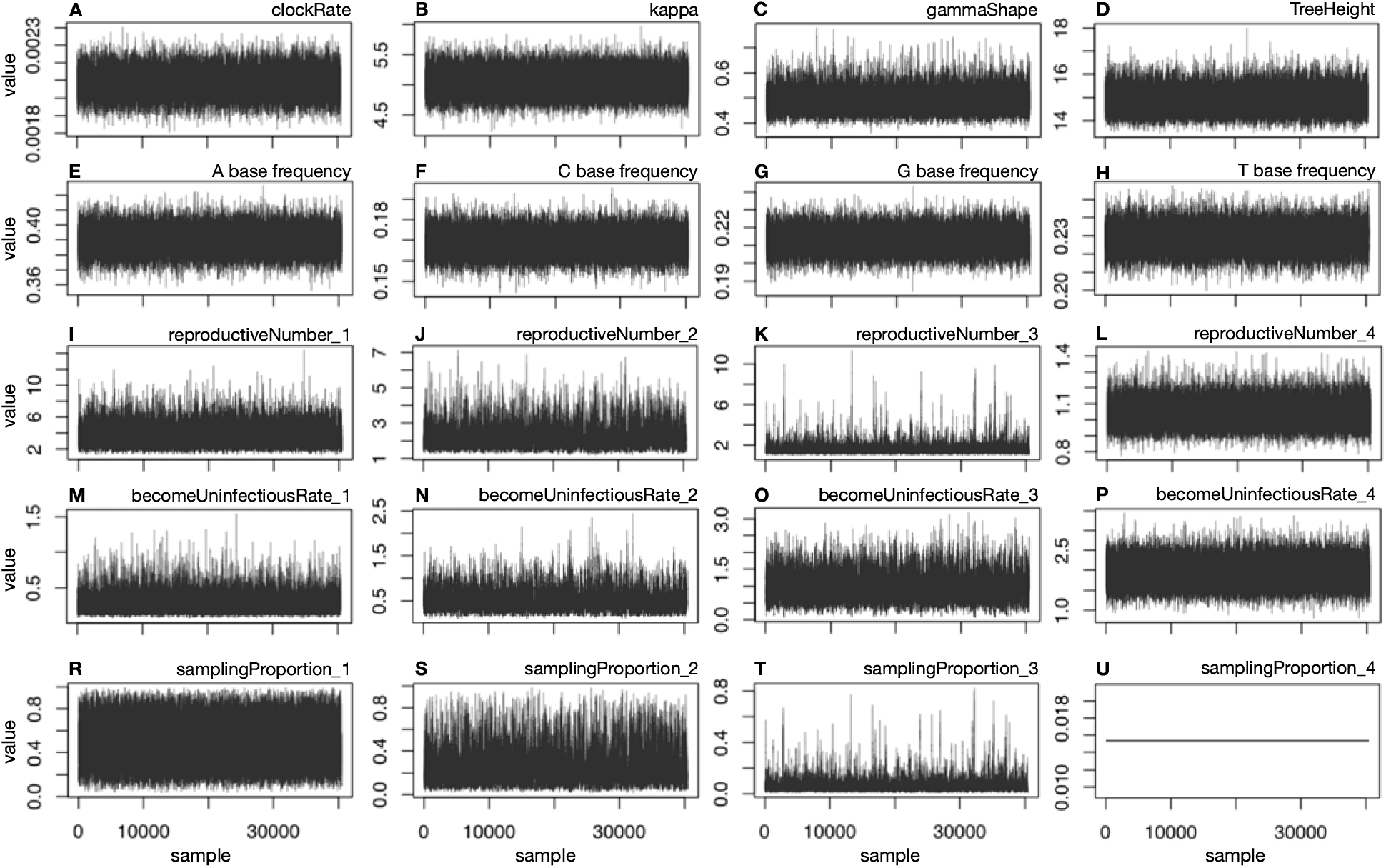
MCMC traces of molecular evolution and epidemiological parameters (BEAST2 run 2_444f). MCMC trace plots of molecular evolution and epidemiological (birth-death-sampling skyline model) parameters generated by BEAST2 (2 independent MCMC chains), based on the sequences simulated under the 2nd hypothetical birth-death-sampling scenario (epidemiological parameters in Supplemental Fig. S7, tree in Supplemental Fig. S3B). Samples are shown after burn-in removal and after thinning of each MCMC chain, and after concatenating the two MCMC chains.

**Figure S17:**
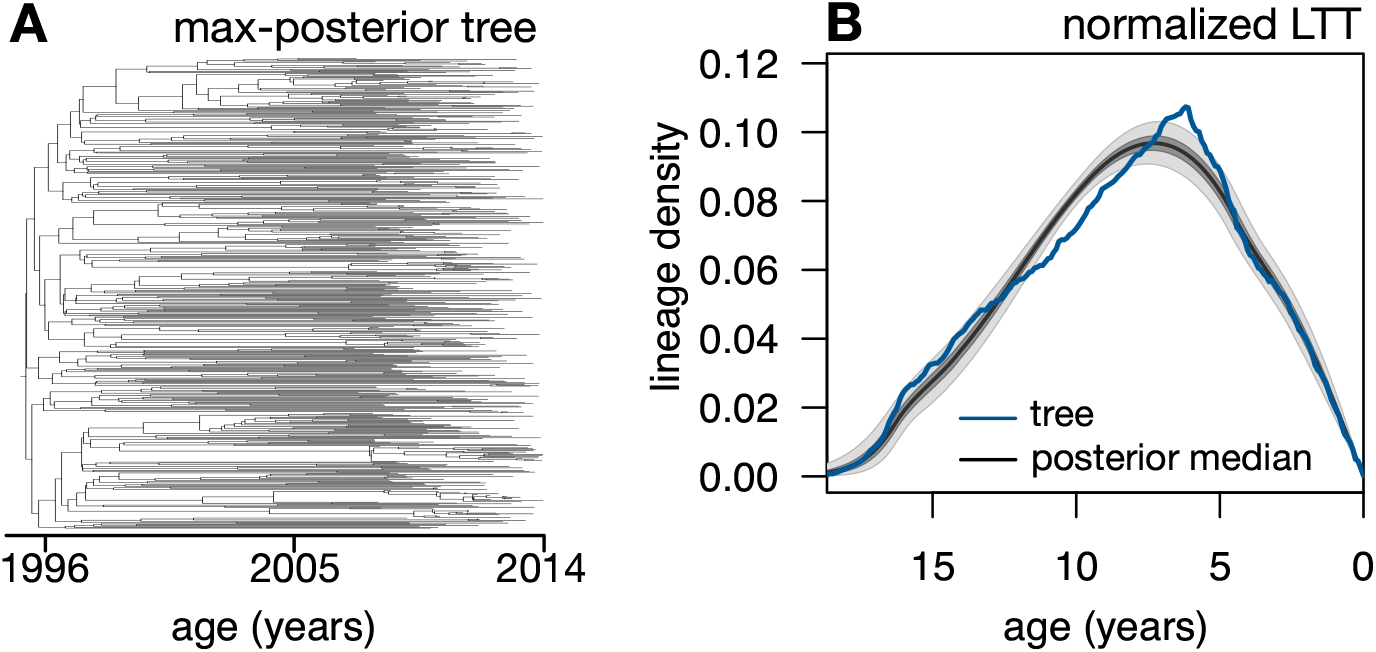
HIV tree and LTT. (A) Maximum-posterior-probability HIV timetree sampled with BEAST2. (B) Lineages-through-time (LTT) curve of the tree normalized to have unit area under-the-curve (blue curve), compared to the normalized deterministic LTTs of the posterior skyline models (black curve shows median, dark and light shades show 50% and 95% equal-tailed credible intervals).

**Figure S18:**
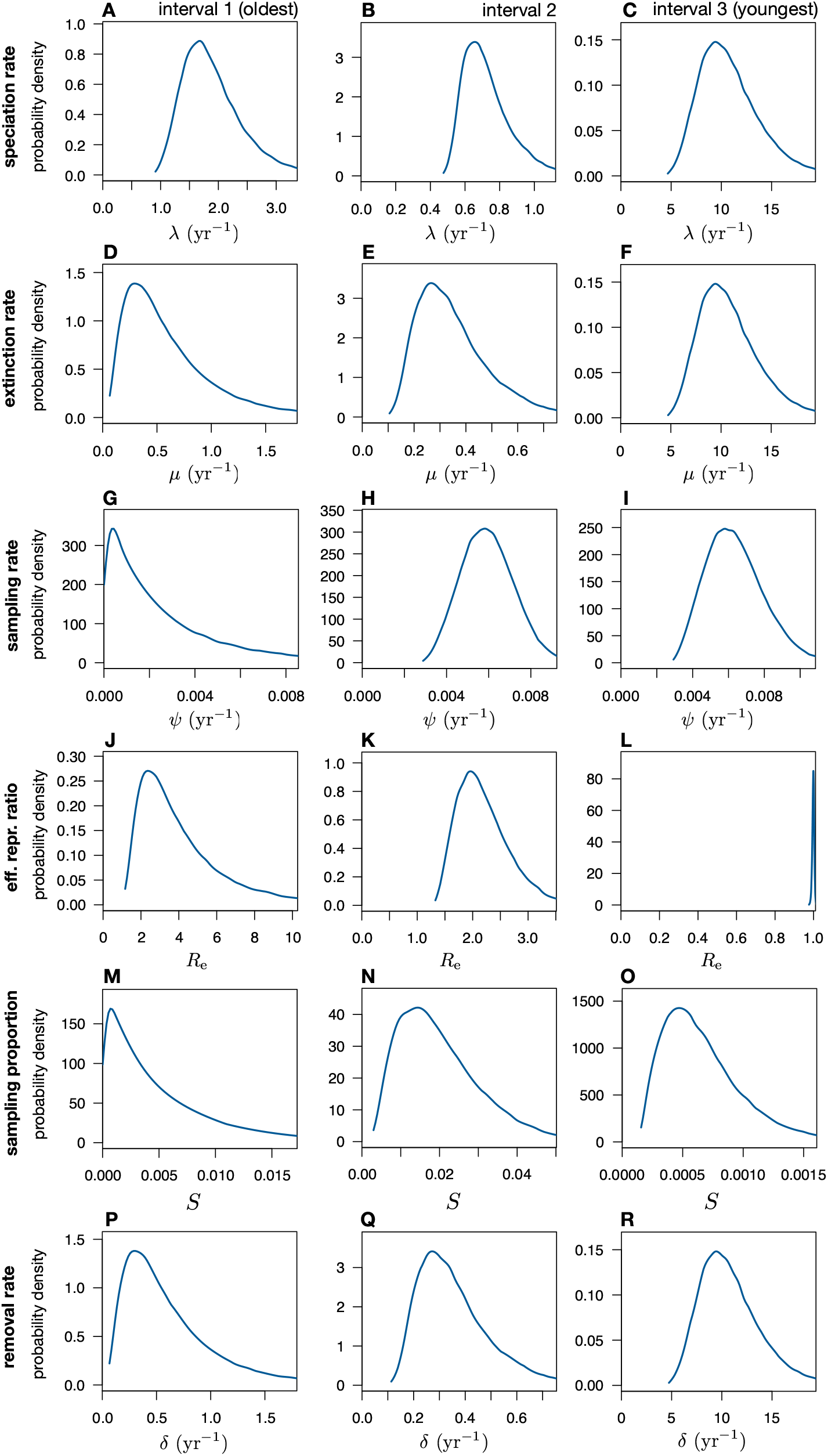
HIV sampled posteriors - probability densities. Probability densities of various epidemiological parameters according to the posterior model distribution sampled by BEAST2, in each of the 3 time intervals. Time intervals cover the periods 0–4, 4–16 and *≥*16 years before present.

**Figure S19:**
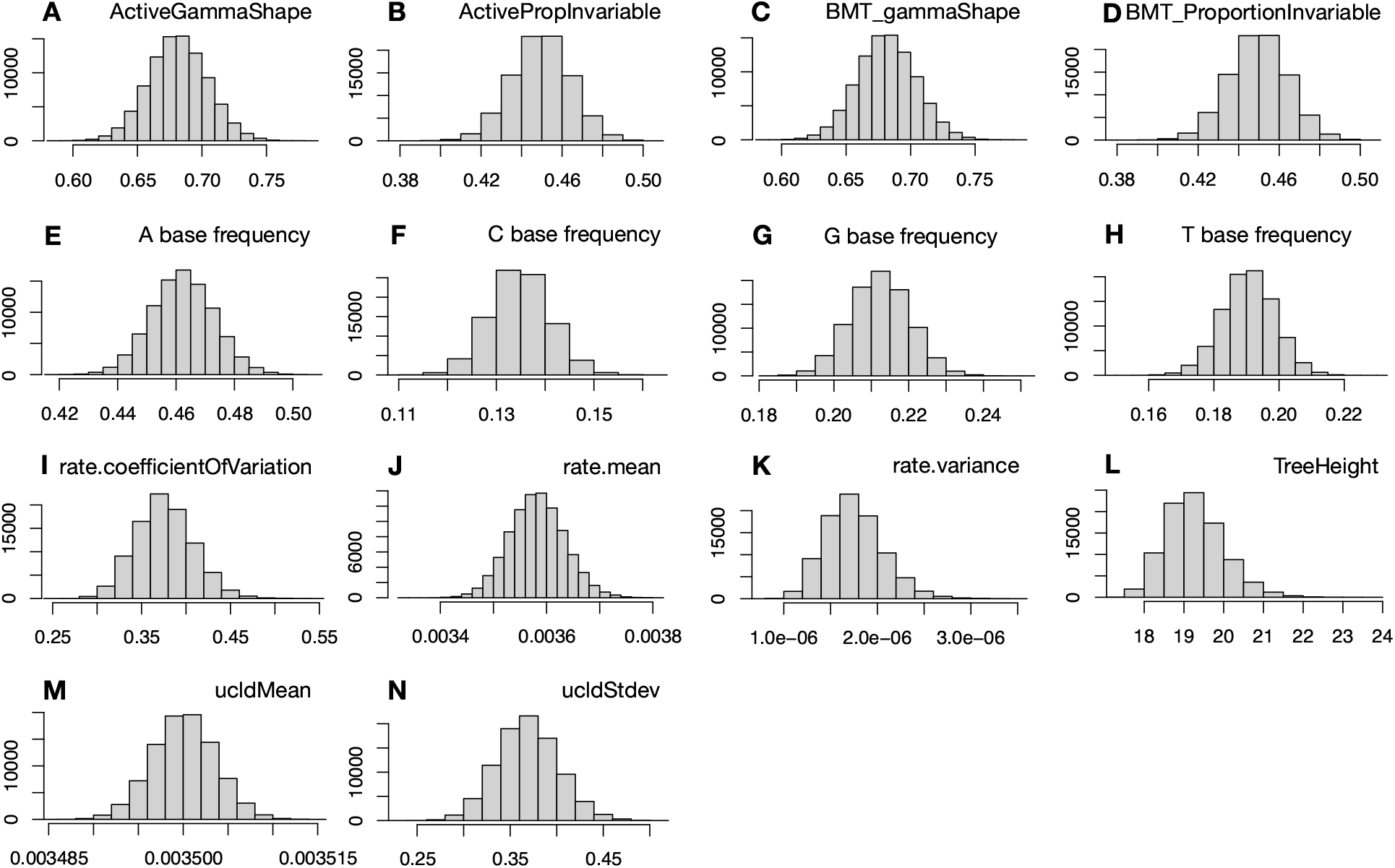
Posterior distributions of molecular evolution parameters (HIV). Histograms of various molecular evolution parameters drawn from the posterior distribution using BEAST2, for the HIV dataset. Vertical bars depict frequencies.

**Figure S20:**
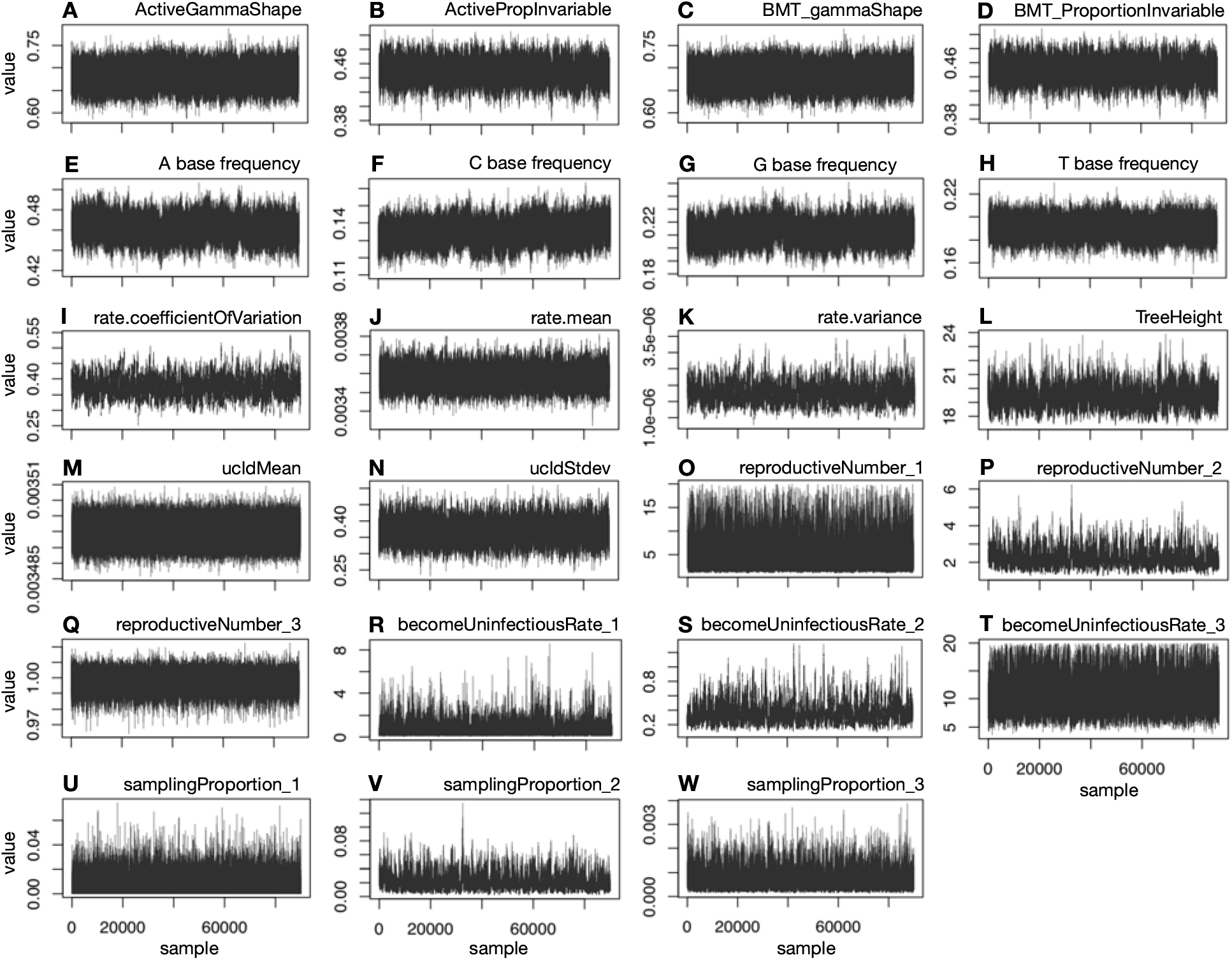
MCMC traces of molecular evolution and epidemiological parameters (HIV). MCMC trace plots of molecular evolution and epidemiological (birth-death-sampling skyline model) parameters generated by BEAST2, for the HIV dataset (2 independent MCMC chains). Samples are shown after burn-in removal and after thinning of each MCMC chain, and after concatenating the two MCMC chains.

**Table S1:** Priors and other settings specified for the BEAST2 analysis of the simulated HIV timetrees. * denotes that the prior parameters are in linear space (default is log).

|  |  |
| --- | --- |
| Tree model | BD skyline serial |
| Substitution model | HKY+ I + G |
| Gamma categories | 4 |
| Clock model | Strict molecular clock |
| Clock rate | LN($\mu=0.003$, $s=1$)* |
| Reproductive number | LN($\mu=0$, $s=1.25$); 4 intervals |
| Become uninfected rate | LN($\mu=2$, $s=1.25$)*; 1 or 4 intervals |
| Sampling proportion | BETA($\alpha=2$, $\beta=2$); 4 intervals |
| Rate change times | 2, 4, 8 years before present |
| Proportion invariant | BETA($\alpha=2$, $\beta=2$) |
| Origin | LN($\mu=20$, $s=1.25$)* |
| Frequency parameter | UNIF(0-1), estimated |
| Gamma shape | EXP(1, offset=0) |
| Kappa | LN($\mu=1$, $s=1.25$) |
| Number of chains | 2 |
| Chain length | 100–200 million |
| Sampling frequency | 10 000 |

**Table S2:** Priors and other settings specified for the BEAST2 analysis of the empirical HIV-1 data from Northern Alberta. * denotes that the prior parameters are in linear space (default is log).

|  |  |
| --- | --- |
| Tree model | BD skyline serial |
| Substitution model | bModelTest, Dirichlet (1, 1, 1, 1) |
| Clock model | Uncorrelated log normal relaxed |
| Clock mean | LN(mu=0.0035, s=0.001)* |
| Clock std.dev. | Gamma(alpha=0.5396, beta=0.3819) |
| Reproductive number | LN(mu=0, s=1.25); 3 intervals |
| Become uninfected rate | LN(mu=0, s=1.25); 3 intervals |
| Sampling proportion | BETA(alpha=1, beta=100); 3 intervals |
| Rate change times | 4, 16 years before most recent sample (2014) |
| Proportion invariant | BETA(alpha=1, beta=4) |
| Origin | LN(mu=25, s=1)* |
| Frequency parameter | UNIF(0-1), estimated |
| Gamma shape | EXP(1, offset=0) |
| Kappa | LN(mu=1, s=1.25) |
| Number of chains | 2 |
| Chain length | 500 million |
| Sampling frequency | 10 000 |

## Notes

### Competing Interest Statement

The authors have declared no competing interest.

